# Systematic analysis of the target recognition and repression by the Pumilio proteins

**DOI:** 10.1101/2024.05.03.592352

**Authors:** Svetlana Farberov, Igor Ulitsky

## Abstract

RNA binding proteins orchestrate the post-transcriptional fate of RNA molecules, but the principles of their action remain poorly understood. Pumilio (PUM) proteins bind 3’UTRs of mRNAs and lead to mRNA decay. To comprehensively map the determinants of recognition of sequences by PUM proteins in cells and to study the binding outcomes, we developed a massively parallel RNA assay that profiled thousands of PUM binding sites in cells undergoing various perturbations or RNA immunoprecipitation. By studying fragments from the *NORAD* long noncoding RNA, we find two features that antagonize repression by PUM proteins – G/C rich sequences, particularly those upstream of the PUM recognition element, and binding of FAM120A, which limits the repression elicited by PUM binding sites. We also find that arrays of PUM sites separated by 8–12 bases offer particularly strong repression and use them to develop a particularly sensitive reporter for PUM repression. In contrast, PUM sites separated by shorter linkers, such as some of those found in *NORAD*, exhibit strong activity interdependence, likely mediated by competition between PUM binding and formation of strong secondary structures. Overall, our findings expand our understanding of the determinants of PUM protein activity in human cells.

**Highlights:**

- A massively parallel assay reports on the binding and activity of Pumilio proteins in human cells
- G/C rich sequence context inhibits repression by Pumilio proteins
- FAM120a binds sequences with Pumilio sites and antagonizes repression by Pumilio proteins
- Arrays of Pumilio binding sites are most effective at distances of 8–12 nt.
- Strong inter-dependency in the tandem Pumilio binding sites in NORAD.

## Introduction

RNA binding proteins (RBPs) shape the post-transcriptional fate of RNA molecules, from transcription to RNA decay (1). Most RBPs have sequence specificity with relatively limited information content that substantially complicates their study. Proteins from the PUF family, which includes the mammalian Pumilio (PUM) proteins PUM1 and PUM2 and other members in diverse eukaryotes from yeast to man, are an interesting exception as they bind specific octamers through a modular structure consisting of eight conserved tandem repeats recognizing one base of RNA each (2).

PUM1 and PUM2 were reported to have largely redundant functions, though some specific activities have been assigned to each of the two proteins (3–5). Both PUM1 and PUM2 (which we will refer to as ‘PUM’ proteins) repress gene expression through recognition of a UGUANAUA binding motif – the Pumilio Recognition Element (PRE) – in the 3′ UTR of mRNAs, likely through recruitment of the CCR4-NOT complex and subsequent degradation of the mRNA target (6). In addition to the canonical and well-established negative effects of PUM proteins, there are reports that they can activate gene expression of some specific genes (7) and globally, through C-rich motifs (8).

The binding sites of PUM proteins on human mRNAs have been mapped by several CLIP-based studies (1, 9, 10). In parallel, the binding preferences of the two proteins were extensively studied in vitro (11–15). These studies mapped the specific details of the PRE, and developed biophysical models for predicting PUM binding to specific sequences. This detailed understanding of PUM biology was also used to design new proteins that can recognize other sequences (16).

The transcripts sensitive to PUM protein levels have also been studied. Both PUM1 and PUM2 mRNAs are PUM targets (3, 17), and so depletion of one protein can be compensated by an increase in the expression of the other. In HEK293 cells, ~1,000 genes are de-repressed when PUM1 and PUM2 are concurrently depleted by siRNA (8) (‘siPUM’). A more recent analysis has specifically measured changes in RNA stability following siPUM (15) and found that ~250 genes were significantly stabilized, whereas 60 genes became significantly less stable. The presence of a PRE is correlated with changes in expression and stability but does not fully explain them, suggesting additional determinants in the sequences of efficient PUM targets (15). Positions closer to the 3’ end of the RNA and A/U content around the PRE have been associated with more efficient repression, though their contributions have not been directly tested yet (15, 18). In addition, there are conflicting reports on the contribution of secondary structure to regulation by PUM proteins (15, 19).

Our lab and the lab of Joshua Mendell identified and characterized the *NORAD* long noncoding RNA (lncRNA) as a cytoplasmic transcript with many PREs that antagonize PUM activity (20–26). The large number of PREs in NORAD, spread across 12 ‘NORAD repeat units’ or NRUs, alongside additional structural features, enable it to overcome its apparent stoichiometric disadvantage over the hundreds of different PUM targets in the cell (25) and to efficiently limit the ability of PUMs to engage with and repress their targets. An increase in NORAD levels leads to the de-repression of PUM target genes, whereas its depletion results in enhanced repression, leading to chromosome instability in NORAD-perturbed human cells and in *NORAD*^−/−^ mice. *NORAD*^−/−^ mice display additional phenotypes such as premature aging (23). Despite at least 18 PREs in the 5.3 kB of *NORAD* RNA sequence, an increase in PUM protein levels has only a limited effect on *NORAD* expression, suggesting that *NORAD* contains elements that protect it from repression by PUM proteins.

Studies of RBPs focused on endogenous RNAs are limited by the high complexity of the gene regulation experienced by individual transcripts, as well as the potential feedback loops they are involved in. Reporter assays have been extensively used to study RBPs, including PUMs (6), but they are limited in the number of different sequences that can be profiled. To address this in other contexts, we and others have been developing and using ‘massively parallel RNA assays’ (MPRNAs), that have been used to study a variety of processes, including splicing (27, 28), translation (29), and RNA export from the nucleus (30–34). MPRNAs involve the design and synthesis of thousands of sequence elements cloned into a plasmid in the context of a reporter gene. The plasmid pool is then transfected into a cell line and reporter RNAs of interest are selected based on their features, such as expression levels, RNA localization, splicing, modification, or translation efficiency. This approach can be easily scaled to determine which sequence variants alter the activities of regulatory elements (31, 32). We recently further extended this approach by combining transfection of an MPRNA library into cells with RNA immunoprecipitation, which allows contrasting the binding of library sequences to a protein of interest with the consequences of the binding, and used this approach to study how HNRNPK protein recognizes and regulates the localization of RNAs containing the SIRLOIN regulatory element (32).

Here, we use a combination of MPRNA and MPRNA-RIP approaches to study the principles of the binding of PUM proteins to their targets and their consequences on mRNA levels in human cells.

## Results

### An MPRNA for dissection of the regulatory actions of PUM proteins

To characterize the sequences that mediate recruitment and regulation by the PUM proteins, we designed a library of 1,825 sequences 140 nt each (PRELibA, **Fig. 1A-B** and **S1A-B**, and **Tables S1-S2**). The library was designed to include PREs in their natural context found in endogenous RNAs, synthetic arrays of PREs to test the effect using particularly sensitive targets with multiple binding sites, and mutants to study the impact of individual bases on PUM repression. Endogenous sequences included sequences from a previous PAR-CLIP study of PUM2 in HEK293 cells (10) and of PUM1 and PUM2 in K562 cells from the ENCODE project (35), as well as PRE-centered fragments of NORAD’s NRUs. We included only endogenous sequences with the UGURNAUA PRE, and added 50 bases upstream and 72 downstream of the motif. The “PRE arrays” were based on the PRE luciferase reporter (36). This reporter has three repeats of a 28 nt sequence containing a UGUACAUA PRE flanked by 14 upstream and 6 downstream bases (**Fig. S1A**). In the library, we systematically shortened these sequences and included PREs separated by 0–20 of the bases in the spacer. We also tried to mutate every possible base to any other bases in this sequence and two shortened versions. We included a corresponding set of mutants for each mutagenesis set where the PRE was mutated to <u>ACA</u>ACAUA (mPREs, included in “mPRE arrays”). We also created arrays from NORAD PREs, which we refer to as N-PREs. In these “N-PRE arrays”, we used the same number of flanking upstream and downstream bases as in the “PRE arrays”. We also included corresponding N-mPRE arrays with UGU→ACA substitutions in the NORAD PREs. Sequences flanking two NORAD PREs (#9 and #10) were mutated systematically, introducing all possible single mutations, which allowed direct comparison with the ‘PRE array’ tiles. For each array, we inserted as many copies as possible within the 140 nt sequence for each of these ‘PRE array’ sequences.

**Figure 1.**
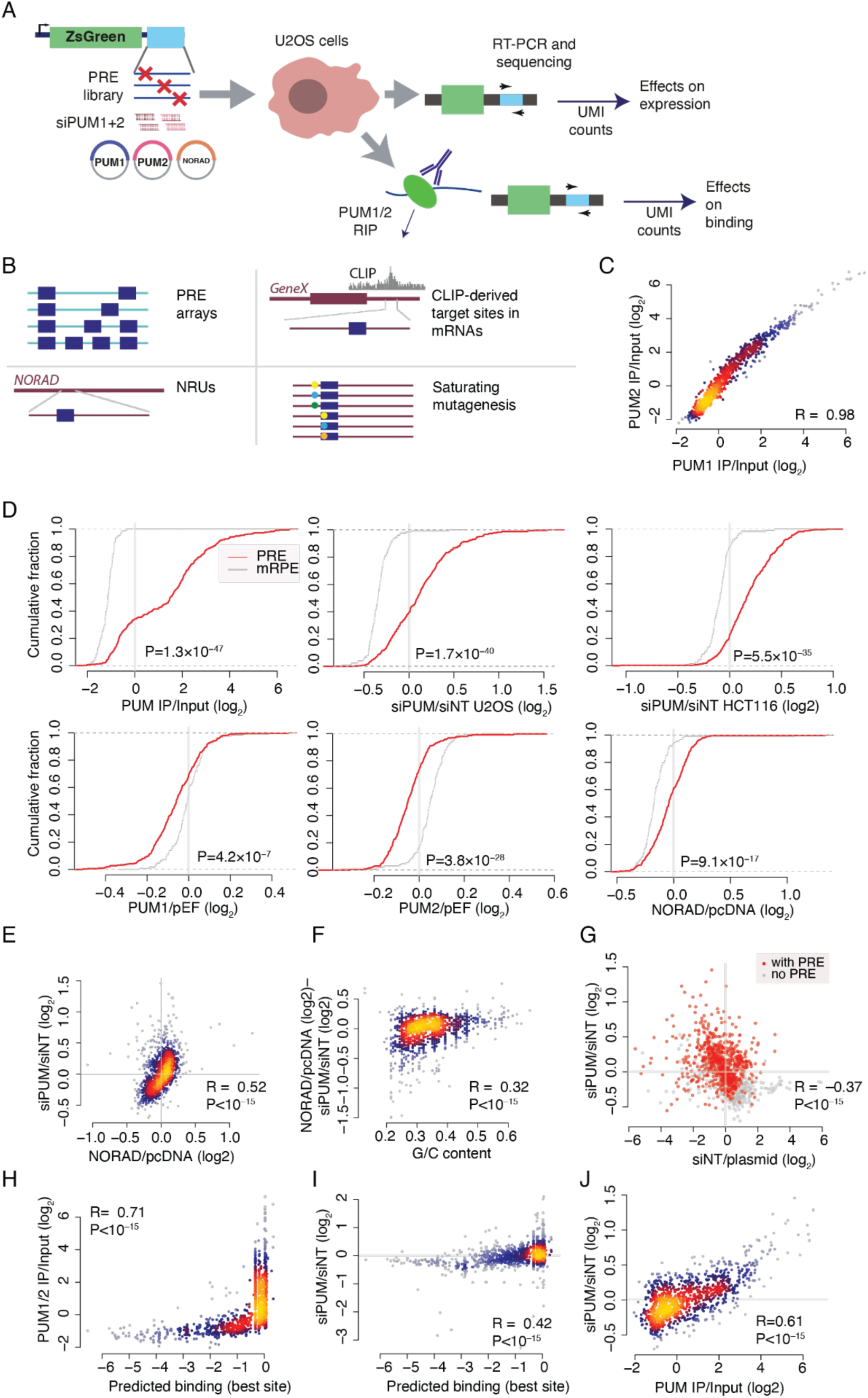
A library for interrogating the functionality and binding of PUM binding sites. **(A)** Outline of the approach. **(B)** Outline of the PRELibA library contents. **(C)** Correlation between the IP/Input ratios of PUM1 and PUM2 MPRNA-RIP experiments. Spearman correlation coefficient is shown. **(D)** Cumulative distribution fraction plots comparing library sequences containing repeated sequences with PREs or mPREs with the indicated ratios. P-values computed using Wilcoxon rank-sum tests. **(E)** Correspondence between response to siPUM treatment and NORAD OE. Color indicates local point density. Spearman’s correlation coefficient is shown. **(F)** as in E comparing the difference between the response to NORAD OE and the response to siPUM as a function of sequence G/C content. **(G)** Correspondence between the expression levels of library sequence expression (normalized to the plasmid) pool and their response to siPUM. Points are colored based on the presence of a PRE in the sequence. **(H)** Same as E, comparing the predicted occupancy of the base with the highest occupancy using the model from (13) and PUM binding in our MPRNA-RIP experiment. **(I)** As in H, comparing the predicted occupancy to the response to siPUM. **(J)** As in E, comparing the enrichment in PUM IP over input to the response to siPUM.

The sequencing tiles were cloned into the 3’ UTR of a zsGreen mRNA and the plasmids were transfected as a pool into U2OS or HCT116 human cell lines which previously (48 h before) underwent various perturbations. Specifically, we knocked down using siRNAs both *PUM1* and *PUM2* mRNAs (‘siPUM’, which reduced the mRNA levels of PUM1 and PUM2 by >60%, **Fig. S1C**) and over-expressed *PUM1*, *PUM2*, or *NORAD* (which over-expressed each transcript by >30-fold, **Fig. S1C**). We then extracted RNA from the cells 24 hr after PRElibA transfection, sequenced the tiles, identified unique molecular identifiers (UMIs, 2.5 million UMIs/library on average), and used biological replicates and DeSeq2 framework (37) to compute the ratios between treated and control cells. Experiments were highly reproducible with Spearman’s R>0.88 between replicates of control cells.

To compare the effects of PUM perturbations with siRNA targeting PUM1 and PUM2 on gene expression vis-a-vis the impact on binding by PUM proteins, we also used MPRNA-RIP (32) with antibodies for PUM1 and PUM2 applied to U2OS cells transfected with PRELibA (**Fig. S1D**). We computed the IP/Input ratio for each sequence that indicates the binding strength. The IP/Input ratios for the two proteins were very similar (Spearman R=0.98, P<10^−15^, **Fig. 1C**), consistent with in vitro studies that found the two proteins to exhibit the same binding preferences (13), and so we combined the data for the two proteins in all subsequence analyses (referring to them as ‘PUM/Input’).

We then examined the effect of siPUM and PUM binding for different groups of sequences, comparing sequences with PREs and mPREs (**Fig. 1D**, **Fig. 2A-C**). As expected, when considering the subsets of sequences with PRE- and mPRE-containing versions, PUM proteins strongly preferred binding the PRE-containing ones (**Fig. 1D**, **2C**). Treatment with siPUM led to de-repression of tiles containing PREs compared to the sets of tiles with mPREs (**Fig. 1D**, **2A-B**). As expected, sequences composed of PRE arrays were more de-repressed than tiles containing individual target sites, either from NORAD or from CLIP targets (**Fig. 2A**). Notably, despite the typically small effect size of PUM perturbations on endogenous transcripts (8, 15), some configurations of PRE arrays led to greater than 2-fold de-repression upon siPUM in our system. A similar hierarchy was observed when considering PUM binding.

**Figure 2.**
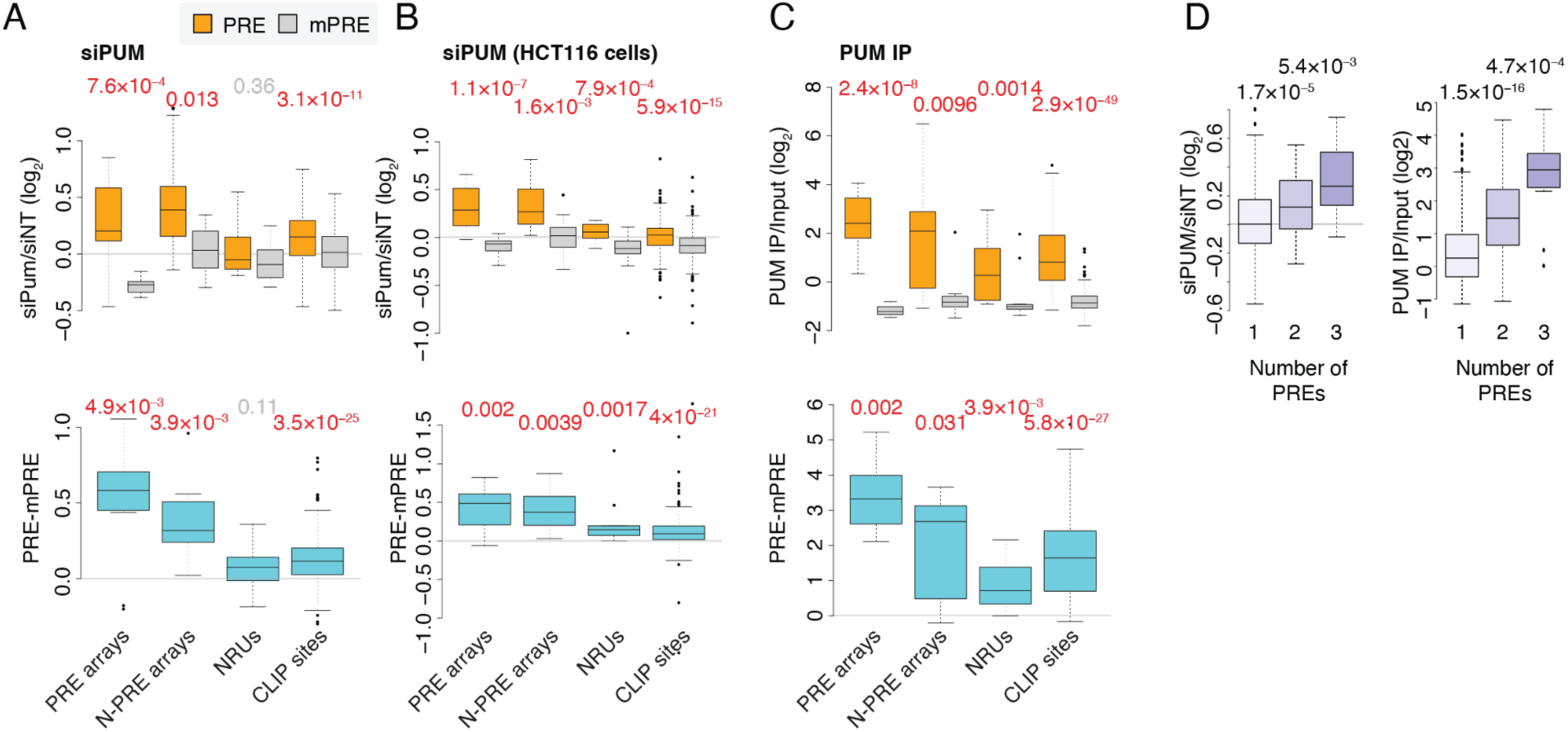
Binding and regulation by PUM of different library subgroups. **(A)** Top: Changes in expression upon siPUM for library sequences in the indicated group in U2OS cells: PRE- or mPRE arrays (**Fig. S1A**); N-PRE- or N-mPRE arrays (‘N-PRE arrays’); sequences derived from NRUs with WT or mutated PREs; PAR-CLIP sites with WT PREs or mutated PREs. Bottom: differences in changes in expression between the WT and corresponding mPRE-containing sequences (i.e., differences between sequences where the only difference is the PRE→mPRE mutation). P-values computed using Wilcoxon rank-sum test. Significant P-values (P<0.05) are in red. Horizontal gray lines are at 0. **(B)** Same as A for HCT116 cells. **(C)** Same as A for PUM binding from the MPRNA-RIP experiment. **(D)** Changes in gene expression following siPUM (left) and changes in binding to PUM (right) for CLIP-derived sequences with the indicated number of PREs.

Given that our library behaves similarly in U2OS and HCT116 cells (**Fig. 1D**, **2A-B**), we proceeded with U2OS cells for the rest of the study. PUM1 or PUM2 overexpression had a milder yet significant effect on repression (**Fig. 1D** and **S1C**), consistent with the repression of PRE-containing transcripts being at near saturation levels in U2OS cells. The high basal PUM activity in U2OS cells makes overexpression less effective; therefore, we focused the rest of the analysis on perturbations that inhibited PUM repression.

When comparing the effects of siPUM and NORAD over-expression (OE, **Fig. S1C**), we observed a milder effect of NORAD OE (median log_2_ fold-change between PRE vs. mPRE containing sequences of 0.03 for NORAD OE vs. 0.11 for siPUM) with a positive correlation of NORAD OE with siPUM (R=0.52, P<10^−15^, **Fig. 1D-E**). Interestingly, some sites that conferred de-repression by siPUM were not substantially affected by NORAD OE. This difference was correlated with the G/C content of the sequences. A/U-rich sequences were preferentially less up-regulated by NORAD OE compared to siPUM (R=0.32, P<10^−15^, **Fig. 1F**), suggesting that NORAD more effectively competes with PUM binding at G/C-rich sites. We focused on the siPUM treatment for the rest of the study, as it conferred the overall strongest effects.

Changes in expression upon siPUM were strongly and inversely correlated with expression levels of the individual tiles, estimated by comparing the RNA sequencing counts with those obtained by sequencing the input plasmids (R=–0.37, **Fig. 1G**), suggesting that repression by PUM explains a substantial portion of the overall variance in the expression of library tiles. Despite the considerable difference in the expression context, and the use of just 140 nt sequence tiles, the siPUM/siNT ratios of CLIP target sites were significantly correlated with the response of their host genes to siPUM in HEK293 cells (8) (Spearman R=0.19, P=2.6×10–5; **Fig. S1E**) and with changes in RNA stability following siPUM (R=0.22 P=2.6×10–5, **Fig. S1E**). PRELibA sequence responses therefore capture the principles of regulation of endogenous transcripts. Furthermore, conservation of the sequence of the PRE element within the tile was significantly, albeit weakly, associated with stronger PUM binding (Spearman R=0.1, P=0.022) and stronger de-repression upon siPUM (Spearman R=0.08, P=0.06), suggesting that PRE that undergo more substantial repression are also more functionally important on the organismal level.

To compare the effects we observe in our PRElibA transfected cells with those shown for PUM binding in vitro, we used the biophysical model from (13) to compute the predicted PUM occupancy on each base in each sequence in the library. This probability was strongly correlated with PUM/Input ratios (R=0.71, P<10^−15^, **Fig. 1H**) and with derepression upon siPUM (R=0.42, **Fig. 1I**).

### Sequence features of effective PUM target sites

We next compared the binding of PUM proteins to library sequences versus the effects of siPUM. We found a strong correlation (Spearman R=0.61, P<10^−15^, **Fig. 1J**), suggesting that sequences better at recruiting PUM proteins are also more effectively regulated by PUMs. We focused further analysis on dissecting the features underlying particularly effective PREs.

First, we examined the collection of WT CLIP sites to evaluate the features associated with effectively repressed and effectively bound sites in diverse contexts. As expected, a larger number of PREs in a tile was associated with stronger de-repression and PUM binding (**Fig. 2D** and **Fig. S1F**). Interestingly, there was a strong influence of the G/C-content of the flanking regions on de-repression of PRE-containing sequences, fitting observations from the analysis of endogenous targets stabilized by siPUM (15). PREs with G/C-rich flanks (G/C>50%) were not de-repressed at all in siPUM (**Fig. 3A**). The G/C-rich flanking regions also led to reduced PUM binding (**Fig. 3B**). To distinguish between the effects on binding and regulation, we stratified the genes based on PUM binding (**Fig. 3C**). In each group with similar binding, siPUM more strongly up-regulated the G/C-poor sequences than the ones with high G/C content, suggesting a stronger effect of G/C content on repression by PUM than on binding (**Fig. 3C**). When considering different PRE sequences, the UGUACAUA variant was significantly less de-repressed than the more common UGUAUAUA and UGUAAAUA PREs (**Fig. 3A**), despite its prevalence, and being the one in a commonly used PRE reporter from the literature. We also considered the C-rich ‘PUM activation’ VYCSWCSC motif previously found in 3’ UTRs of transcripts down-regulated upon PUM perturbation (8). Consistently with the endogenous targets, sequences bearing this motif were down-regulated upon siPUM (**Fig. 3A**). However, there was no evidence that PUM proteins bound this motif (**Fig. 3B**).

**Figure 3.**
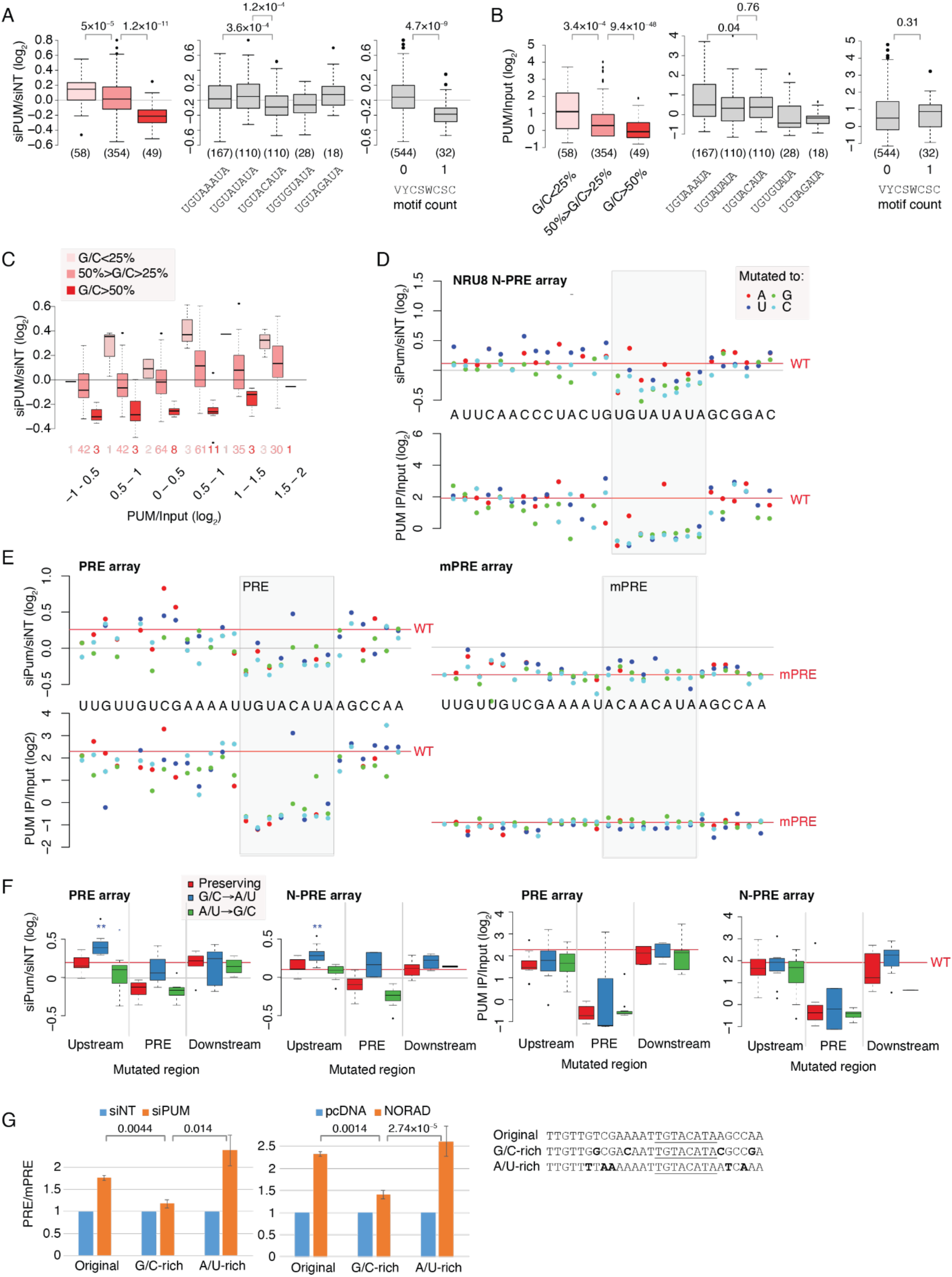
Increased G/C content in regions flanking the PRE affects PUM binding and regulation. **(A-B)** siPUM regulation (A) and PUM binding (B), considering only sequences with one PRE, changes in expression and binding for the indicated overall G/C content of the sequence, indicated PRE sequence, and presence of the VYCSWCSC motif (V=A/C/G, Y=C/U, S=G/C, W=A/T). Horizontal gray line is at 0. P-values were computed using a two-sided Wilcoxon rank-sum test. Numbers in parentheses indicate the number of sequences in each group. **(C)** CLIP-derived sequences were divided into groups based on their PUM/Input ratios and, within each group, further separated by their G/C content. The response to siPUM is compared between the groups. Horizontal gray line is at 0. **(D)** Changes in gene expression upon siPUM (top) and in binding to PUM (bottom) for single-nucleotide mutants of the NORAD repeat unit (NRU) 8-derived PRE-surrounding sequence. Each mutated sequence was repeated for a total length of 140 nt. The red lines indicate the levels of the WT sequences with no mutations. Horizontal gray line is at 0. The PRE-containing region is shaded in gray. **(E)** As in D, for mutations in the PRE reporter-derived sequence (left) or the mPRE-derived sequence (right). **(F)** Changes in expression upon siPUM (left) and binding to PUM (right) for the indicated mutated sequence, grouping together positions upstream (13 bases) of the PRE, within the PRE, and downstream of the PRE (6 bases). * - P<0.05, ** - P<0.005, Wilcoxon rank-sum test, comparing to the ‘preserving’ mutations. **(G)** Luciferase level ratios between plasmids containing three PRE or mPRE arrays, either the original from (36) or recoded to have A/U- or G/C-flanking regions, responding to siPUM (left) or NORAD OE (right) treatments, in each case, the ratio is normalized to the control. P-values computed using two-sided t-test.

To determine the contribution of single bases to repression and binding, we examined the arrays of mutated sequences, testing individual mutations (**Fig. 3D-E**), and aggregating mutations in the different regions by their effect on the G/C content (**Fig. 3F**). As expected, mutations in the consensus bases of the PRE strongly affected both PUM binding and derepression upon siPUM treatment (**Fig. 3D-E**). Mutations that decreased G/C content in the flanking regions, in particular, increased de-repression upon siPUM treatment, especially when the more G/C-rich NORAD sequence from NRU8 was mutated. Some of these mutations also affect PUM binding, but overall, G/C→A/T mutations make these sequences more sensitive to PUM without significantly affecting binding. Consensus PREs flanked by a G/C-rich context, such as those found in NORAD, are thus effective at binding PUMs while eliciting relatively limited repression.

To test these findings in an independent system, we replaced the PREs in the PRE reporter that we based our PRE arrays on (36) with more G/C-rich or A/U-rich sequences (**Fig. 3G**). We transfected these new reporters alongside the original into U2OS cells transfected with siNT, siPUM, pcDNA3.1 (control plasmid) or pcDNA3.1-NORAD. As expected, PUM inhibition or NORAD over-expression de-repressed the reporter (**Fig. 3G**). The G/C-rich reporter was significantly less de-repressed than the original reporter or the A/U-rich reporter, fitting with the results obtained through the PRELibA analysis, and showing that PREs flanked by A/U-rich are indeed more potently repressed (**Fig. 3G**).

### Optimal spacing between PREs facilitates an improved reporter for PUM activity

One of the challenges with studying PUMs is the limited effect size that their repression elicits on the targets with a single PRE. The reporter developed by Goldstrohm lab and used by us and others (8, 20, 26) contains three UGUACAUA PREs, but its effect is still limited, as siPUM causes a ~75% de-repression. A subset of the library containing PRE repeats separated by 0– 20 bases allowed us to systematically compare the efficiency of dense PRE arrays. We note that the sequences differed in the number of PREs and the spacing between them (**Fig. S1A**). When examining de-repression (siPUM/siNT ratios), we found that PREs separated by 8–12 bases (which had 7–9 PREs) underwent the most robust repression, associated with more potent PUM binding, though the effects on repression where more prominent than the effects on binding (**Fig. 4A-B**). Surprisingly, these sequences were more repressed than those with 10 or more PREs and separated by less than 7 bases. A possible explanation for this observation is that the PRE sequence has a relatively strong tendency to pair with one another. As a result, sequences with short inter-PRE distances have relatively very stable predicted structures, more so than those separated by 8–12 bases (**Fig. 4C**). The sequence separated by 8–12 bases thus gave the best tradeoff between a large number of PREs and a relatively loose structure.

**Figure 4.**
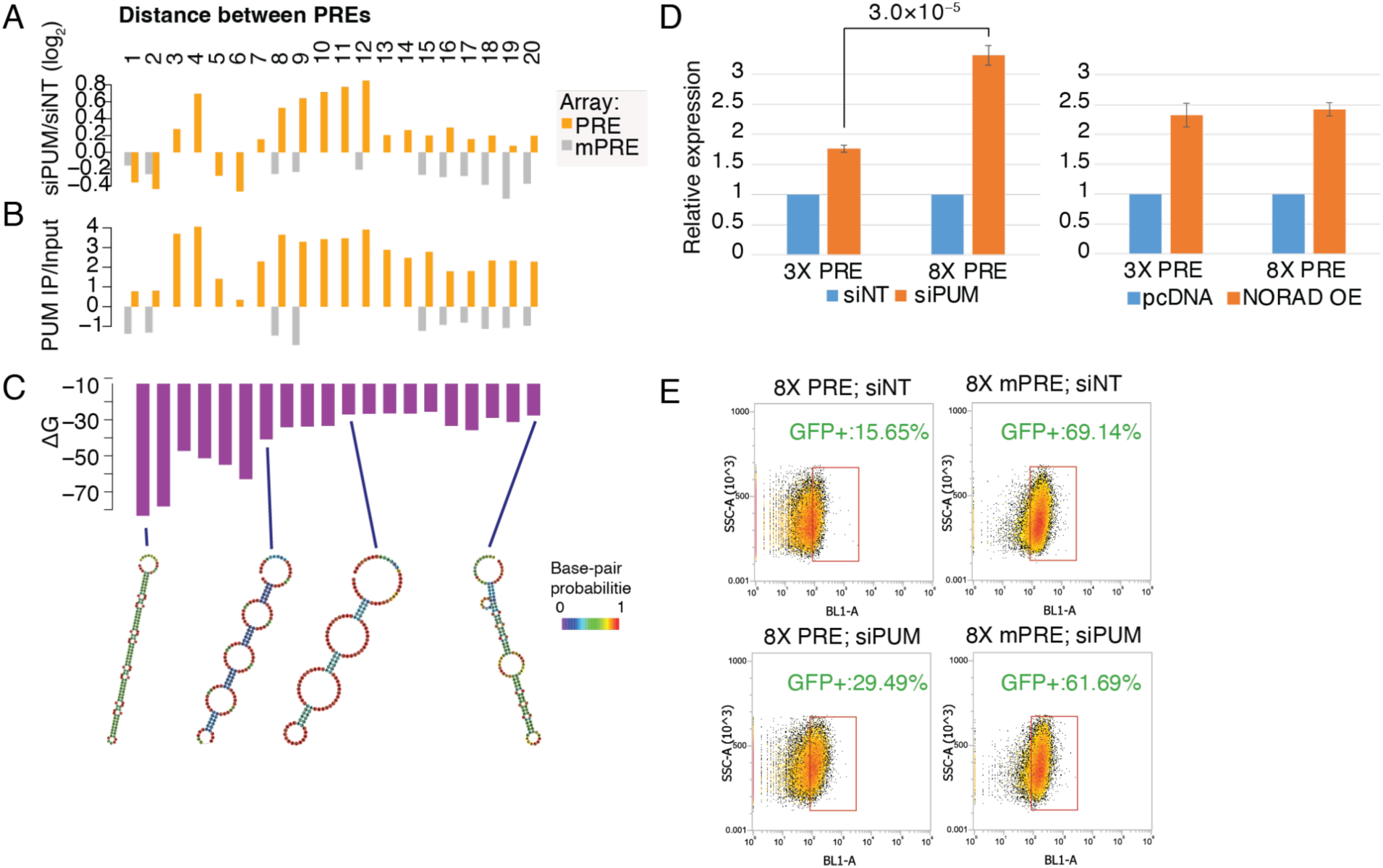
Optimal spacing between PREs. **(A)** Changes in expression upon siPUM for PREs or mPREs separated by the indicated number of bases from the PRE reporter (**Fig. S1A**). **(B)** As in A, for PUM binding. **(C)** Predicted secondary structure minimal free energy for each sequence in A–B. Selected predicted structures are shown at the bottom with bases color-coded based on the base-pair probabilities. **(D)** Luciferase reporter results comparing the indicated reporters with 3X original PREs vs. 8X PREs separated by 12 base spaces AAUCAAGAAAAU. Values are ratios between PRE and mPRE reporters normalized to the controls (siNT or pcDNA3.1 empty vector). P-value computed using two-sided Student’s t-test. **(E)** FACS plots comparing cells with an integrated GFP reporter having 8X PREs or an 8X mPRE array in the 3’ UTR, treated with siNT control or with siPUM.

Based on these results, we designed and synthesized a new reporter, containing 8 repeats of UGUAUAUA separated by 12 A/U-rich bases (<u>UGUAUAUA</u>AAUCAAGAAAAU). Indeed, this reporter responded much more strongly when tested in cells transfected with siPUM. Interestingly, the response to NORAD OE remained unchanged (**Fig. 4D**), consistently with the relatively lower efficacy of NORAD competing with A/U-rich sequences in PRELibA (**Fig. 1F**). Furthermore, integration of this reporter into the 3’ UTR of an mRNA encoding an AcGFP fluorescent protein resulted in a visible difference in fluorescence upon siPUM treatment, which could be quantified by FACS (**Fig. 4E**). The PRELibA-based design thus allows the development of a sensitive assay for quantification of PUM activity.

### Limited effects of predicted secondary structure on PUM binding and repression

The observation that multi-PRE arrays with extensive structure had a limited response to siPUM, led us to examine more closely the relationship between the predicted RNA structure and binding/regulation by PUM proteins. We hence focused on the 471 PRELibA sequences that had exactly one PRE and predicted their minimal energy secondary structures using RNAfold (38). As expected from the results of the effects of mutations on G/C content, sequences with stronger predicted folds were significantly less likely to be up-regulated by siPUM, and, to a lesser extent, were less likely to be bound by PUM proteins (**Fig. 5A**). However, the correlations with the strength of the predicted structure were lower than correlations with G/C content, when considering the same sequence set (**Fig. 5B**). Therefore, it is difficult to disentangle the effects of PRE accessibility that can be affected by secondary structure from the potential sequence-specific binding of other RBPs to G/C rich sequences (see Discussion).

**Figure 5.**
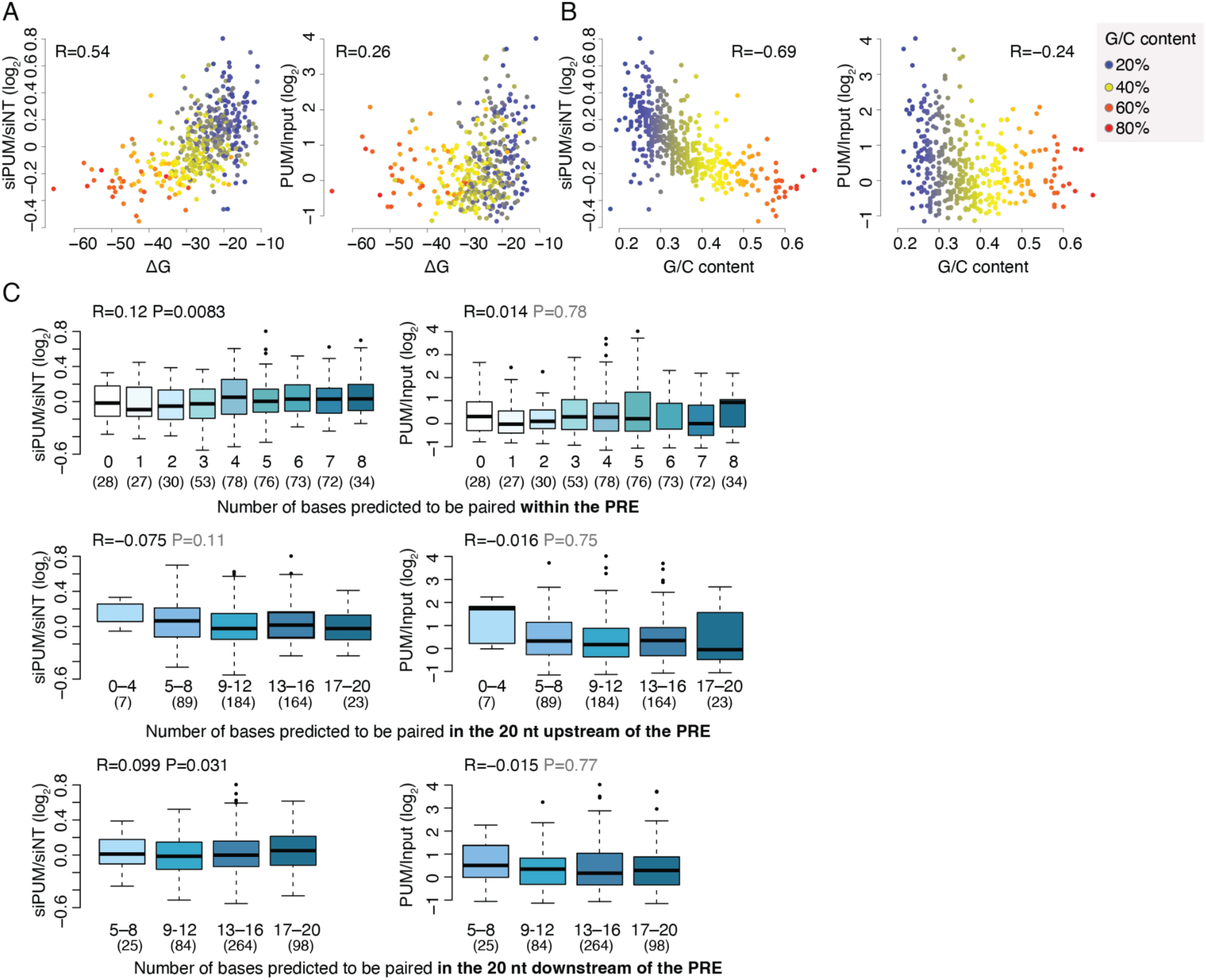
Relationship between predicted RNA structure and PUM proteins. **(A)** Correlations between the stability of the minimal free energy structure and changes in gene expression following siPUM (left) and binding by PUM proteins (right). Spearman correlations and P-values are indicated. **(B)** Same as A, for the G/C content of the library sequence. **(C)** Changes in gene expression following siPUM (left) and binding by PUM proteins (right), for sequences with the indicated number of paired bases in the indicated region. Numbers of sequences with the indicated structure are in parentheses below the number of paired bases.

We next examined the specific minimal free energy structure predicted by RNAfold around the single PRE in each sequence. Surprisingly, bases predicted to be paired within the PRE motifs did not appear to have adverse effects on PUM binding or regulation by PUM. In fact, there was a weak yet significant positive correlation between the number of paired PRE bases and with de-repression by siPUM (**Fig. 5C**). When considering the 20 bases upstream or downstream of the PRE, the correlations were also weak, with a notable increase in de-repression upon siPUM for particularly unstructured regions upstream of the PRE (<8/20 paired bases). This observation is consistent with the mutagenesis results (**Fig. 3E-F**), which were performed on different sequences, and where G/C→A/U mutations, which would generally weaken the secondary structure, led to higher de-repression specifically when occurring upstream of the PRE.

These results are consistent with previous *in vitro* studies, that observed largely no correlation between PUM binding and predicted secondary structure (13) and suggest that either PUM binding is associated with helicase activity that can unwind structures effectively to expose the PREs, or that the accuracy of the specific predicted folds is low. In any case, it appears that particularly structured regions affect PUM binding, possibly by making the PRE inaccessibly, yet small structures around the PRE have a more limited effect, perhaps restricted to the region immediately upstream of the PRE.

### FAM120A binds sequences enriched with PREs and antagonizes PUM activity

FAM120A (aka OSSA/C9ORF10) is a relatively poorly studied cytoplasmic protein recently shown to antagonize microRNA-mediated repression by associating with Ago2 and binding G-rich motifs (39). We previously found that FAM120A binds NORAD near PUM binding sites (26). FAM120A was also enriched in the IP of NORAD fragments, and its eCLIP peaks over NORAD overlap with those of PUM1 and PUM2. Out of all the RBPs in the ENCODE eCLIP data FAM120A peaks preferentially overlapped those of PUM2, and this overlap was one of the most substantial across all the RBPs profiled by ENCODE (**Fig. 6A**). Notably, PUM1 eCLIP peaks did not tend to overlap with FAM120A (**Fig. 6A**). We hypothesized that FAM120A is associated with PUM-bound sequences and modulates repression by PUMs.

**Figure 6.**
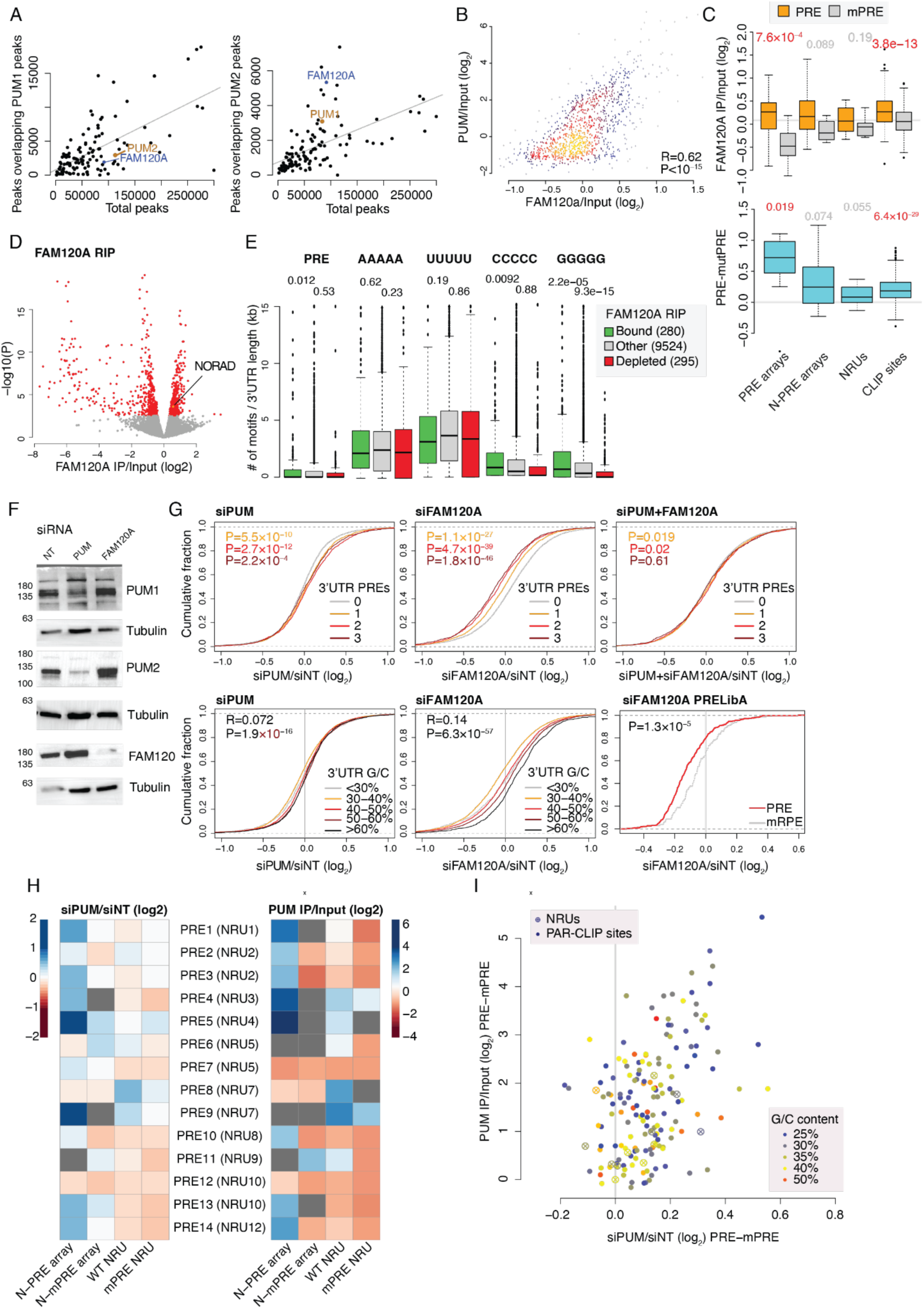
FAM120A is recruited to PREs and antagonizes PUM repression. **(A)** Correspondence between the total number of eCLIP peaks for all the factors profiled by ENCODE and the number of peaks that overlap the peaks of PUM1 (left) or PUM2 (right). **(B)** Correspondence between binding of FAM120A and PUM in the MPRNA-RIP experiments. The color indicates local point density. Spearman correlation coefficient and P-value are shown. **(C)** As in Fig. 2, a comparison between the indicated values for groups of sequences containing PREs (orange) or mPREs (gray). The ratios are shown in the top plots, and the differences between the corresponding PRE- and mPRE-containing sequences are shown at the button. P-values are computed using Wilcoxon rank-sum tests and significant (P<0.05) P-values are in red. **(D)** Volcano plot for the enrichment in FAM120A RIP sample (x-axis) and the significance of the enrichment (y-axis) as computed by DESeq2 for endogenous RNAs annotated in GENCODE v42 in U2OS cells (n=6 biological replicates). Genes with adjusted P<0.05 are in red and *NORAD* is in orange. **(E)** Number of 3’ UTR sequence matches for the indicated motifs, normalized per 1Kb of 3’ UTR length, in the 3’ UTRs of genes enriched or depleted in the FAM120A RIP sample compared to the input (adjusted P<0.05 and enrichment/depletion of at least 50%). P-values are shown for comparisons between the bound or the depleted and the other 3’ UTRs, computed using two-sided Student’s t-test. **(F)** Western blots for the indicated factors in U2OS cells treated with the indicated siRNAs. **(G)** Cumulative distribution plots comparing the indicated groups of 3’UTRs or PRELibA sequences. P-values computed using two-sided Wilcoxon rank-sum test. **(H)** Derepression by siPUM (left) and binding to PUM (right) for arrays of NORAD-derived PREs or mPRE controls or fragments from NRUs and their mPRE controls. Dark gray squares represent missing values. **(I)** Correspondence between derepression by siPUM and PUM binding for PAR-CLIP–derived PUM binding sites and NRU sequences, color-coded by the G/C content of the tile.

To characterize the effects of the binding of FAM120A to the PRELibA sequences, we immunoprecipitated FAM120A (**Fig. S2A**) and performed MPRNA-RIP in the same conditions as those used for PUM. We also sequenced endogenous RNAs bound by FAM120A in the same RIP samples. FAM120A binding to PRELibA sequences strongly correlated with PUM binding (R=0.62, **Fig. 6B**), with no notable difference when considering PUM1 or PUM2 (R=0.65 and R=0.61, respectively), and FAM120A preferentially bound sequences containing PRE elements over other sequences (**Fig. 6C**). FAM120A also preferentially bound endogenous RNAs containing PREs, including NORAD, consistently with the ENCODE eCLIP data (**Fig. 6D-E**). Within PRELibA, FAM120A preferentially bound A/U-rich sequences, more so than PUM (**Fig. S2B**). 3’UTRs of endogenous FAM120A targets were enriched with PREs and G- and C-homopolymers (**Fig. 6E**). Consistently with FAM120A being recruited to PREs, there was a very strong correlation between FAM120A binding and the response to siPUM (R=0.65). Mutations in the PRE sequence preferentially abolished FAM120A binding (**Fig. S2C**), suggesting that PREs specifically recruited FAM120A, directly or indirectly.

We then knocked down FAM120A in U2OS cells (‘siFAM120A’ **Fig. 6F** and **S2D**) and examined changes in expression of endogenous RNAs and PRELibA sequences. Knockdown of FAM120A down-regulated transcripts containing PREs in their 3’ UTRs and sequences in PRELibA containing PRE. The effect on the endogenous transcripts was lost when both PUMs and FAM120A were depleted (**Fig. 6G**), consistently with a protective effect of FAM120A against PUM-mediated decay. When considering different subsets of sequences in PRELibA, FAM120A depletion down-regulated PRE arrays and NORAD sequences (**Fig. S2E**), but only the changes in NORAD sequences were significant compared to the effects on mPRE sequences. Among the PRE arrays, the effect of the KD was strongest on the closely spaced PREs (**Fig. S2F**). Interestingly, KD of FAM120A increased PUM protein levels (**Fig. 6F**), suggesting that beyond shared binding sites, FAM120A may directly affect PUM proteins, and this up-regulation can contribute to the down-regulation of PUM targets upon FAM120A KD. We conclude that FAM120A is recruited to PRE-containing sequences that are generally responsive to siPUM and limits their repression, which is enhanced when FAM120A is lost.

### PUM binding and potential regulation of *NORAD* repeats

PRELibA contains sequences from NORAD, including N-PRE arrays – bases from 14 different *NORAD* PREs with flanking bases (and corresponding N-mPRE arrays) and other sequences with longer context around these PREs (not repeated, which we refer to as ‘NRUs’). As expected, N-PRE arrays that strongly bound PUM proteins were also typically up-regulated by siPUM. Exceptions were N-PREs where the PRE was UGUAAAGA, UGUGUAUA, and UGUAUGUA (PRE7, PRE8, and PRE12 in **Fig. 6H**) – two of which diverged from the PRE consensus. As expected, N-PRE arrays, that contain multiple PREs, were more regulated than NRUs that contain just one or two PREs (**Fig. 6H**). Overall, the distributions of PUM binding and derepression by siPUM were similar between *NORAD*-derived sequences and those derived from PAR-CLIP sites. *NORAD*-derived fragments were relatively weak PUM targets, but not weaker than expected based on their relative GC-richness (**Fig. 6I**). Among the *NORAD* PREs, the most robust PUM binding was to NRU 7, which had a tandem pair of PREs, and which is part of the sequence required for the minimal functional version of *NORAD* that we recently characterized (21). We conclude that most of the PREs within *NORAD* are functional in binding PUM proteins, and at least some of them can be regulated by PUM proteins when expressed in isolation outside of the context of the full-length *NORAD*.

### Co-dependent closely spaced PREs in *NORAD*

PRELibA contained only a few mutated sequences. To get a broader view of the sequence determinants regulating PUM binding, we designed a second library – PRELibB – with 6,294 oligos 140 nt each (**Table S3**). In PRELibB we included more extensive mutagenesis of eight sequences, including two NORAD fragments from NRUs 3 and 7, four PUM2 CLIP sites from PAR-CLIP data (10) (from *HIAT1*, *MGEA5*, *PGBD4*, and *AP1S1* mRNAs), and two target sites from PUM1 eCLIP data (in *SMARCA2* and *IGF2BP1* mRNAs). The selected sequences represented different levels of binding and different response levels to PUM perturbation in the PRELibA data.

For each of the eight core sequences in PRELibB, each containing at least one PRE, we mutagenized both the WT and a sequence where each PRE was mutated with UGU→ACA. In addition to all single point mutations, we also mutated consecutive 4mers, 6mers, or 10mers, using mutations designed to maintain G/C content (A⇔U and G⇔C). Importantly, the mutated sequences differed in the number of PREs, with two tandem PREs in NRUs 3 and 7, a pair of more distant PREs in PGBD4, and a single prominent PRE in MGEA5, AP1S1, and IRF2BP1. PRELibB also included repeats of 20mers containing a PRE or a mutated PRE separated by 12 bases with all possible mutations.

For seven of the nine mutated sequences, we observed the expected behavior where some of the mutations decreased the ability to bind and to be regulated by PUM, with high correlations between the effects (**Fig. 7A** and **Fig. S3**). In contrast, in sequences that lacked a functional PRE, the mutations had limited effects. For SMARCA2, selected as an example of a sequence repressed by siPUM in PRELibA, both WT and mutated sequences were poorly bound by PUMs, though mutating the PRE still had some effect (**Fig. S3**).

**Figure 7.**
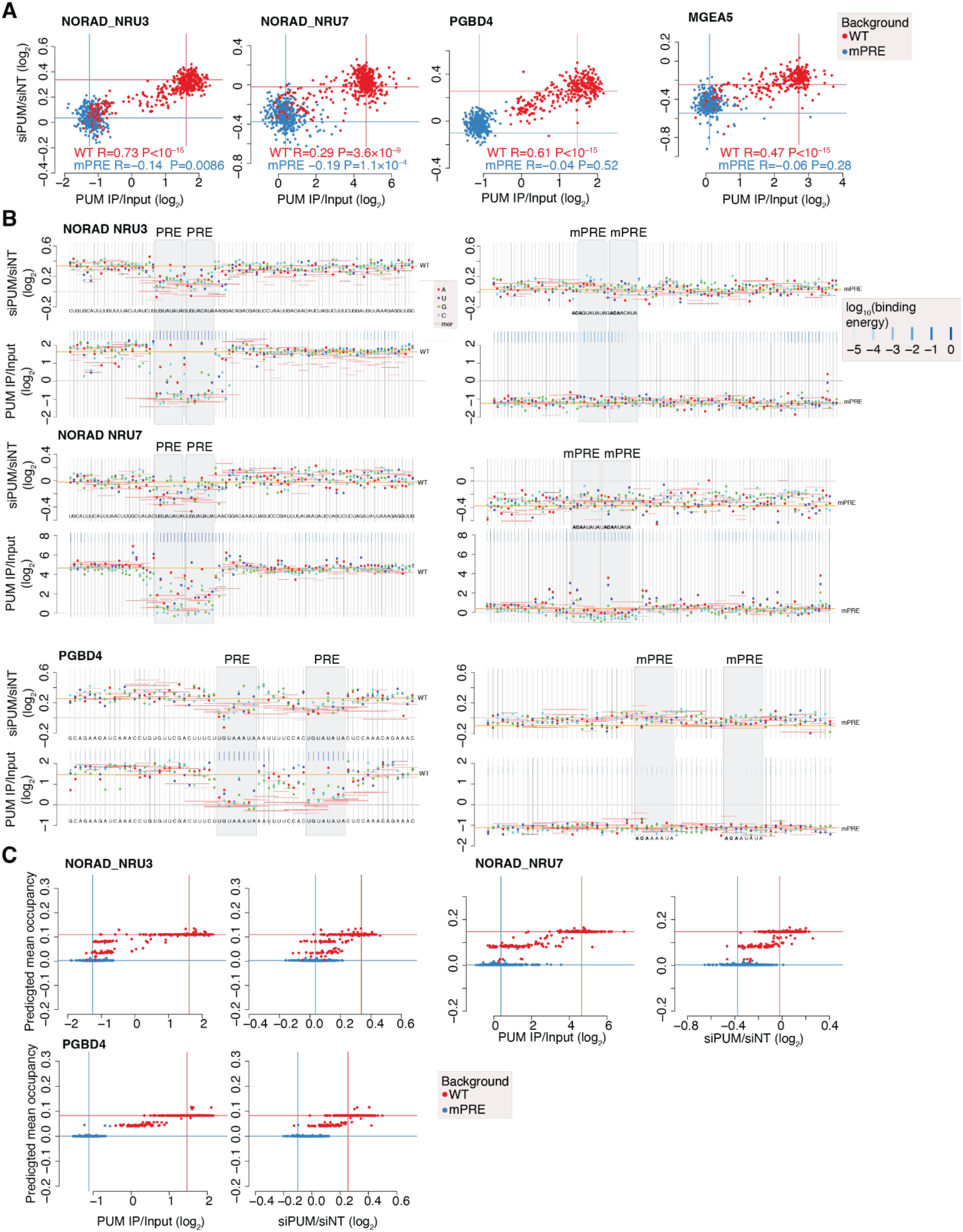
PRELibB analysis. **(A)** Correspondence between PUM binding and siPUM response for sequence variants of WT or PRE-mutated sequences. Four of the sequences in PRELibB are shown; the others are in **Fig. S3**. Lines indicate the ratios for the baseline WT and mPRE sequences. Spearman correlations and P-values are shown separately for WT and mPRE sequence mutants. **(B)** Changes in siPUM response (top) and in PUM binding (bottom) for the indicated single-nucleotide variants (dots) and kmer-mutants (short vertical lines covering the mutated positions). Long orange horizontal lines indicate the ratios for the WT or mPRE baseline sequences. Blue vertical lines are colored based on the predicted occupancy of PUM on each base (see legend). Additional sequences are shown in **Fig. S3**. **(C)** Correlation between PUM binding or response to siPUM and the mean predicted occupancy of PUM on the mutated sequence for variants of the WT or the mPRE sequences.

For *HIAT1*, the mutations had a limited effect on PUM binding and regulation, while as expected, sequences with an intact PRE expressed significantly better in siPUM cells (Wilcoxon two-sided rank=sum test P<10^−15^, **Fig. S3D**). The *HIAT1* sequence was particularly A/U-rich and contained some near-PRE motifs, where mutations that brought them closer to PRE consensus improved binding, suggesting that these sites could potentially compensate even for the loss of all canonical PREs in this sequence. Supporting this notion, there was a very strong correlation between predicted mean PUM occupancy based on the *in vitro* model and binding to and regulation by PUMs (R=0.38 for PUM/Input and R=0.29 for siPUM/siNT ratios), suggesting that the *HIAT1* sequence is mostly regulated directly by PUM binding.

We next examined the interdependence between pairs of PREs. Strikingly, in the sequences from NRUs 3 and 7, mutations in either of the two adjacent PREs had a very strong effect on PUM binding, bringing the PUM/Input ratios close to that of a sequence where both sites were mutated (**Fig. 7B**). Similarly, regulation by PUM was affected to a similar degree when either site or both were mutated (**Fig. 7B**). In contrast, in *PGBD4*, where the PREs were separated by 10 nt, mutations in each PRE affected both binding and repression, but less than the double mutant, giving rise to an intermediate effect on both binding and repression (**Fig. 7B**). Similarly, when examining the predicted occupancy of PUM on the mutated sequence, mutations in either of the PGBD4 PRE led to intermediate predicted occupancy and intermediate binding/repression, whereas mutations in the NORAD NRU PREs completely abolished both binding and repression, while only an intermediate reduction in occupancy was predicted by the biophysical model (**Fig. 7C**). The tandem PREs in NRUs are thus a powerful PUM recruitment module (as evident in **Fig. 6H**, where PRE9 from NRU7 is the strongest responder), but exhibit an unusual inter-dependence between the closely spaced PREs separated by just one base.

*MGEA5*, *AP1S1*, *IRF2BP1* contained a single PRE and mutations affecting both PUM binding and response to PUM KD clustered within this PRE (**Fig. S3A-C**). In *AP1S1* there were some additional UGU-containing sequences quite far from the PRE consensus where mutations affected binding substantially. Interestingly, in *AP1S1* mutations, increasing A/U content generally positively affected both binding and repression.

For the sequence from *SMARCA2*, which was more G/C rich, the UGUGGAUA PRE did not appear functional, and the only mutations that substantially improved binding were kmer mutations that converted A/G-rich sequence stretches to T/C-rich ones (**Fig. S3E**). Point mutations had a limited effect on the expression of SMARCA2-derived sequences upon PUM KD, arguing against this repression being driven by a specific sequence motif. This supported the notion that the repression of sequences having C-rich motifs upon PUM KD is not due to a particular motif recognized by PUM proteins but is indirect.

## Discussion

Our study complements previous efforts that interrogated the function of PUM proteins. Previous studies used CLIP to study PUM binding to endogenous RNAs (10, 22, 40). CLIP has the advantage of being unbiased by prior knowledge and thus allows the identification of binding sites that do not necessarily contain canonical PREs. However, CLIP-based studies are limited in the numbers and nature of the different sequences interrogated, and endogenous transcripts vary widely in their features beyond PREs. Other studies used RNA sequencing to examine the consequences of perturbation of PUM proteins on gene expression and mRNA stability (8, 15, 22, 41, 42). Our MPRNA-based approach has several advantages, as we can compare thousands of carefully designed sequences, systematically mutate individual positions, and study arrays of short sequences in which small effects can be effectively amplified and interrogated. By combining MPRNAs and MPRNA-RIPs, we can also compare side-by-side binding and repression by PUM proteins of different sequences, all of which are transcribed from the same promoter and have the same sequence backbone, eliminating much of the variability between endogenous PRE-bearing 3’ UTRs. We could examine the effects of sequence changes on both binding and function, which allows us, for example, to directly test the previously predicted contribution of A/U-richness to PUM regulation (15).

There are also some important limitations to our approach. By design, to limit variability, we aimed to place most of the PREs in the same relative position within the library sequences, which also means that the PREs are mostly found at the same distance from the end of the coding sequence and from the poly(A) tail, in the context of relatively short 3’UTRs. Endogenous PREs are typically found in longer 3’ UTRs, and the position within the 3’ UTR was suggested to influence PUM repression (15), with more effective PREs found closer to the poly(A) tail in a rend resembling effective miRNA binding sites (43). Future studies can potentially place the library in different host contexts or positions within a longer 3’ UTR, akin to our recent approach to study context-specific effects on mRNA localization (30). Another limitation of our PRE array approach is that when we changed the spacers between the PREs in PRE arrays, we trimmed a single uniform spacer sequence (derived from a baseline PRE reporter). In future studies, it will be interesting to try and test multiple sequences, particularly non-uniform spacers, that will also influence the secondary structure of the PRE arrays in different ways. While this manuscript was in preparation, Hass et al. described a related approach they termed MPRNA-IP for massively parallel interrogation of binding by PUM proteins (44). Their library design is very different from ours in that they selected a few PUM targets and tiled across the entire length of the mRNA rather than focusing on a larger number of CLIP-derived and synthetic target sites.

Our libraries included a series of fragments from the NORAD sequence. NORAD harbors an unusually large number of PREs, yet its levels exhibit relatively mild sensitivity to changes in PUM activity (20, 22). In our MPRNA, the responses of NORAD-derived sequences appear to be similar to those of other sequences - dense arrays of NORAD-derived PREs typically bind PUM proteins well and respond strongly to siPUM (**Fig. 6H**). Individual NRU-derived sequences are more G/C-rich than some of the CLIP-derived PUM targets. So, their repression is relatively mild (**Fig. 6H**), which is explained at least in part by the G/C bases flanking the PRE (**Fig. 3D**).

FAM120A, which we identify here as a PUM antagonist, is another potential candidate for stabilizing NORAD despite PUM binding. NORAD has multiple FAM120A binding sites in the ENCODE CLIP data (26) and it is one of the endogenous RNAs most recovered by FAM120A RIP (**Fig. 6D**). In the MPRNA, individual NRUs didn’t appear to show particularly strong FAM120A binding beyond that of other PUM targets (**Fig. 6D**). Still, it is possible that the collective binding of multiple NRUs to FAM120A in the context of the full-length NORAD, alongside the antagonistic effect of FAM120A towards PUM repression, helps maintain NORAD stability while it is bound to PUM. If that is indeed the case, FAM120A-mediated stabilization co-operates with the relative G/C-richness of the NRUs, and the sequestration of NORAD within NORAD-PUM phase-separated bodies (25), which may explain why NORAD is not significantly down-regulated by FAM120A knockdown.

Several aspects of regulation by PUM proteins, including the additive effects of multiple binding sites and the importance of A/U-rich flanking regions, resemble those of microRNAs, which is plausible since both primarily act by recruiting the CCR-NOT complex that triggers mRNA deadenylation. The nature of the contribution of the flanking A/U-richness remains incompletely understood in both cases. A/U-rich sequences form less stable secondary structures than G/C-rich ones, but it is not yet clear if this is the main or only explanation for the observed phenomena. For miRNAs, earlier studies found that site accessibility does not contribute to repression when A/U-richness is taken into account (43). Still, more complex models built on larger datasets did find that site accessibility also has independent, yet apparently weaker contribution (45). It remains possible that A/U-richness acts via site accessibility but the latter is hard to estimate, or that A/U-richness is important because of various protein binding preferences rather than its effect on PRE accessibility. A recent study found that FAM120A inhibits miRNA-mediated repression (39). That study implicated G-rich sequences as the drivers of FAM120A recruitment, although A- and U-rich homopolymers were also found to bind FAM120A (39, 46). In our hands, we primarily see FAM120A binding both A/U-rich PRE-bearing sequences, and G- and C-rich homopolymers. Future studies will elucidate the specific nature of the elements recognized by FAM120A and how it inhibits both PUMs and microRNA effectors. A compelling hypothesis is that the shared inhibition of miRNAs and PUMs is related to inhibition of the interaction with CCR-NOT, or that FAM120A binding is relocating the target mRNAs to areas in the cell where PUMs and microRNAs are less active.

We had a limited ability to look at cooperativity between PUM binding sites, since in our PRE arrays, both the spacing between the PREs and the structure of the whole sequence are affected. Since PREs match their reverse complement sequences quite well, dense arrays of PREs have a high propensity to form hairpins (**Fig. 4C**). Such hairpins in NRUs potentially lead to bimodal PRE behavior - if both PREs are intact, PUM proteins effectively bind the sequence, but if one is mutated, the other PRE loses its potency. Surprisingly, even larger perturbations of one of the PREs in the tandem pair, such as changes of four bases, that would be expected to open the structure substantially have a strong effect. Interestingly, the tandem PREs in NORAD are highly conserved in evolution (**Fig. S4**), with three such tandem pairs in the human sequence and two in the mouse found in four different odd-number NRUs (3, 5, 7, and 9). The single-base spacing between the tandem PREs is also highly conserved, although the base itself differs in the different NRU pairs.

We find overall no evidence for any differences between PUM1 and PUM2 in binding or in the consequences of their over-expression, which is consistent with the *in vitro* massively parallel measurements (13, 15) and surprising given the phenotypes ascribed to the loss-of-function of *PUM1* and *PUM2*, which are not identical, and quite severe (3–5). Possible explanations for this apparent discrepancy are that one of the proteins or both can have distinct interaction partners that allow them to act independently of RNA binding, or that their levels in cells, that are sometimes different (22, 47), lead to differential dependencies on their accumulation. In any case, we conclude that PUM proteins act on the same target sites and have similar consequences to their binding.

In addition to the PREs, perturbation of PUM proteins was previously reported to lead to down-regulation of mRNAs carrying a C-rich motif (8, 15). In our system, we replicate PUM-mediated repression of transcripts containing such motifs but see no evidence that PUM proteins bind this motif, consistent with an in vitro study that also failed to observe such binding (13), suggesting that regulation of these motifs is robust across multiple cellular models, but is most likely indirect.

Our systematic approach for elucidating the principles of binding and activity of PUM proteins can be directly applied to other RBPs. As we show, the combination of MPRNAs and MPRNA-RIP experiments is particularly powerful, as is the combination of sequences bearing individual binding motifs in endogenous RNAs with designed arrays of binding sites and their mutants – binding site arrays allow to amplify effects of specific mutants, and the endogenous sequences allow comparison of large numbers of contexts surrounding individual sites. As PRELibB exemplifies, secondary libraries built around specific endogenous sequences with interesting behaviors can further delineate their functional determinants. Future MPRNA-based studies will majorly contribute to elucidating the binding-function axes of additional RBPs and deciphering an important layer of gene regulation.

## Materials and Methods

### PRELibA design and cloning

Sequences of the PRE arrays were based on the PRE sequence from the PUM reporter (36) UUGUUGUCGAAAAU<u>UGUACAUA</u>AGCCAA (PRE underlined) in which first the 1–14 of the bases upstream of the PRE, and then the 6 bases downstream of the PREs were iteratively deleted. Each sequence was repeated as many times as possible to the full length of 140 nt (**Fig. S1A**). The same procedure was applied to the mutated UUGUUGUCGAAAAU<u>ACAACAUA</u>AGCCAA sequence (mPRE arrays). Systematic mutagenesis was applied to the 26 nt PRE array sequence, as well as the shortened AU<u>UGUACAUA</u>AGCCAA and <u>UGUACAUA</u> arrayed sequences. The repeated sequences from NORAD are in **Table S2**. In addition to arrays of these N-PREs, the arrays of the PREs UAUAC<u>UGUAUAUA</u>U<u>UGUAUAUA</u>UAACGG and AUUCAACCCUAC<u>UGUGUAUAUA</u>GCGGAC were systematically mutated. For the CLIP studies, peaks with a score of at least 3 were considered, and only those with the UGURNAUA motif. In each peak, the 50 nt upstream of the first PRE was included, followed by the PRE and another 82 bases to the full length of 140 nt.

Designed oligo pools were synthesized by Twist Biosciences (San Francisco, CA). Oligo pools were amplified by PCR. PCR products were purified and inserted into the 3’ UTR of a zsGreen mRNA in the pZsProSensor-1 vector by restriction-free cloning and transformed into E. cloni classic electrocompetent cells (Lucigen, 60117-1). Plasmid DNA was isolated and representation of the pool was verified by Illumina sequencing.

### PRELibB design and cloning

Baseline sequences were selected from PRELibA (**Table S3** for baseline sequences and numbers of variants and **Table S4** for library sequences). Bases or kmers in positions 20–120 in WT or mPRE baseline sequences were systematically mutated. In *HIAT1*, *MGEA5, PGBD4*, and *AP1S1* PAR-CLIP–based sequences (WT and mPRE), positions 10–80 were mutated and kmers in positions 1–90 were perturbed. In the ENCODE eCLIP-derived *SMARCA2* sequence, bases and kmers in positions 10–100 were mutated. In the ENCODE eCLIP-derived *IRF2BP1* sequence, bases and kmers in positions 8–98 were mutated. Mutated windows differed between the baseline sequences to fit the final library size of <6,000 sequences.

### Tissue culture

Human osteosarcoma cell line U2OS (ATCC-HTB-96) and human colorectal carcinoma cell line HCT116 (ATCC-CCL-247) were cultured in DMEM (Gibco, 11-965-092) supplemented with 10% fetal bovine serum (FBS) and 100 U of penicillin/0.1 mg mL−1 streptomycin at 37°C in a humidified incubator with 5% CO2.

### Plasmid and siRNA transfections

Transfections of plasmids were performed using GenJet in Vitro DNA Transfection Reagent (SignaGen Laboratories) for U2OS cells and polyethyleneimine (PEI) for HCT116 cells. PUM proteins were overexpressed by transfecting cells with pEF-Flag-PUM1 and PUM2 plasmids from (48); pEF-BOS empty vector was used as a control. *NORAD* was overexpressed using a plasmid containing its full sequence in the pcDNA3.1 backbone (Addgene #120383, from (20)); pcDNA3.1 (+) (Invitrogen) was used as a control. In transfection, 400 ng of plasmid was used per 200,000 cells in 6-well plates, the cells were transfected with PRElib plasmid 48 hr post PUM / NORAD transfection.

*NORAD*, *PUM1*, *PUM2*, and *FAM120A* were knocked down using siRNA pools at a final concentration of 25 nM (all from Dharmacon, pools of four siRNAs per target, **Table S5**); mammalian non-targeting siRNA (Lincode Non-targeting Pool, Dharmacon) was used as control. PRElib plasmid was transfected 48 hr post siRNA transfection.

### MPRNA

U2OS and HCT116 cells were transfected with PRELibA or B, as described above, 48 hr post overexpression or knockdown. Total RNA was isolated 24 hr post transfection using TRI Reagent (MRC). RNA samples were reverse transcribed and amplified by PCR using primers containing unique sequencing identifiers as well as the sequences required for illumina sequencing.

### MPRNA-RIP

RNA Immunoprecipitation (RIP) from whole-cell extracts (49) was performed as follows. Protein A/G magnetic beads (GeneScript) were washed in NT2 buffer (50 mM Tris HCl pH 7.4, 150 mM NaCl, 1 mM MgCl_2_, 0.05% NP-40) and conjugated with antibodies against PUM1 (5 ug, Bethyl, A300-201A), PUM2 (5 ug, Abcam, ab10361), FAM120A (5 ug, Abcam, ab156695), or Rabbit IgG as a negative control at room temperature for 1 hr in gentle rotation.

U2OS cells were collected 48 hr after library transfection (2.5 µg of DNA per 5×10^6^ cells). Cells were lysed for 5 min on ice in Polysome lysis buffer (100 mM KCl, 5 mM MgCl_2_, 10 mM HEPES NaOH pH 7, 0.5% NP-40, 1 mM DTT, protease inhibitor cocktail (ApexBio), RNase inhibitor (EURx, E4210)) and centrifuged at 20,000 g for 10 min at 4^0^C. Part of the supernatants was saved as total cell lysate input. The rest, containing 2-3 mg protein extract, was incubated for 3 hr at 4^0^C in gentle rotation with antibody-bound beads. The beads were washed six times with a lysis buffer, each time separated by magnetic force. The remaining mixture of magnetic beads-antibodies-protein-RNA complexes was divided in two – half was mixed with laemmli buffer and boiled at 95^0^C for 5 min for further analysis by Western blot while the other half was incubated with Proteinase K for 30 min at 55^0^C with gentle shaking to remove proteins. The remaining RNA was extracted by TRI Reagent. RNAs isolated from the IP were reverse transcribed and amplified by PCR to generate sequencing libraries. Western blot was used to verify the precipitation of the desired proteins.

### MPRNA data analysis

Sequencing reads (at least 150nt in length) were processed and aligned to the library sequences using the MPRNA data processing (https://github.com/IgorUlitsky/MPRNA) also used in previous studies (30, 32), unique molecular identifiers (UMIs) associated with each sequence in each sample were counted. Libraries were also prepared from the plasmids containing the library, and 199 sequences that had less than 30 reads in the plasmid library were excluded from further analysis. DeSeq2 (37) was used for computing log2-transformed fold changes, and those were only used for tiles for which DeSeq2 computed an adjusted p-value, to exclude tiles with too low or too variable expression. Secondary structures were predicted for the library sequences using RNAfold (38). Occupancies of PUM based on the *in vitro* model from (13) were predicted using the script from https://github.com/pufmodel/Pumilio_occupancy_predictions.

### qPCR analysis of gene expression

Extraction of total RNA for KD or overexpression verification was done 24 hr post PRElib transfection using TRI Reagent. RNA was reverse transcribed using the qScript Flex cDNA kit (Quanta bioscience) with a mix of oligo dT and random primers according to the manufacturer’s instructions. Gene expression was determined using the ViiA7 qPCR with the Fast SYBR qPCR mix (Life Technologies). The primer sets used for the different genes are listed in **Table S6**. All gene expression levels are represented relative to their relevant control (ΔCt) and normalized to GAPDH (ΔΔCt).

### Luciferase assays

Luciferase PRE reporters were generated by inserting PRE repeats into the 3’UTR of Renilla luciferase in the psiCheck-2 dual luciferase reporter vector (Promega) (**Table S7**). The reporters co-transfected with siRNA pools (siNT or siPUM) using LipoJet siRNA/DNA co-transfection reagent (SignaGen) or with NORAD or pcDNA3.1 plasmids usingGenJet (SignaGen). Luciferase activity was determined 48 hr post transfection using the Dual-Glo Luciferase Assay System (Promega) in the Microplate Luminometer device (Veritas). A relative response ratio, from Renilla and Firefly luminescence signals was calculated for each sample and values were normalized to the controls.

### Generation of reporter cell lines

U2OS reporter cell lines expressing AcGFP fused to an arrays of 8X PRE and mPRE in its 3’UTR were generated by targeting the construct into AAVS1 locus using the CRISPR–Cas9 system. Targeting vector was generated by replacing the OsTIR1 sequence in pMGS46 plasmid (Addgene, #126580, from (50)) with AcGFP-PRE/mPRE. The puromycin resistance cassette was replaced by hygromycin resistance. The reporters were co transfected with a plasmid carrying the Cas9 gene and a guide RNA for the human AAVS1 locus (pMGS7 Addgene, #126582). Cells were cultured at low density under hygromycin selection (100 µg/ml) and single clones were selected.

Reporter cell lines were treated with siNT or siPUM. Cells were harvested 48 hrs post transfection and GFP expression levels were determined by FACS (Attune™ NxT Flow Cytometer, Life). Data was analyzed using the Attune cytometric software.

## Statistical analysis

Unless indicated otherwise, all pairwise comparisons were done using the Wilcoxon rank-sam test, and all correlations were computed using Spearman correlation.

## Data availability

All the sequencing data has been deposited in the SRA database under the BioProject accession PRJNA1028573.

## Supp Figures

**Figure S1.**
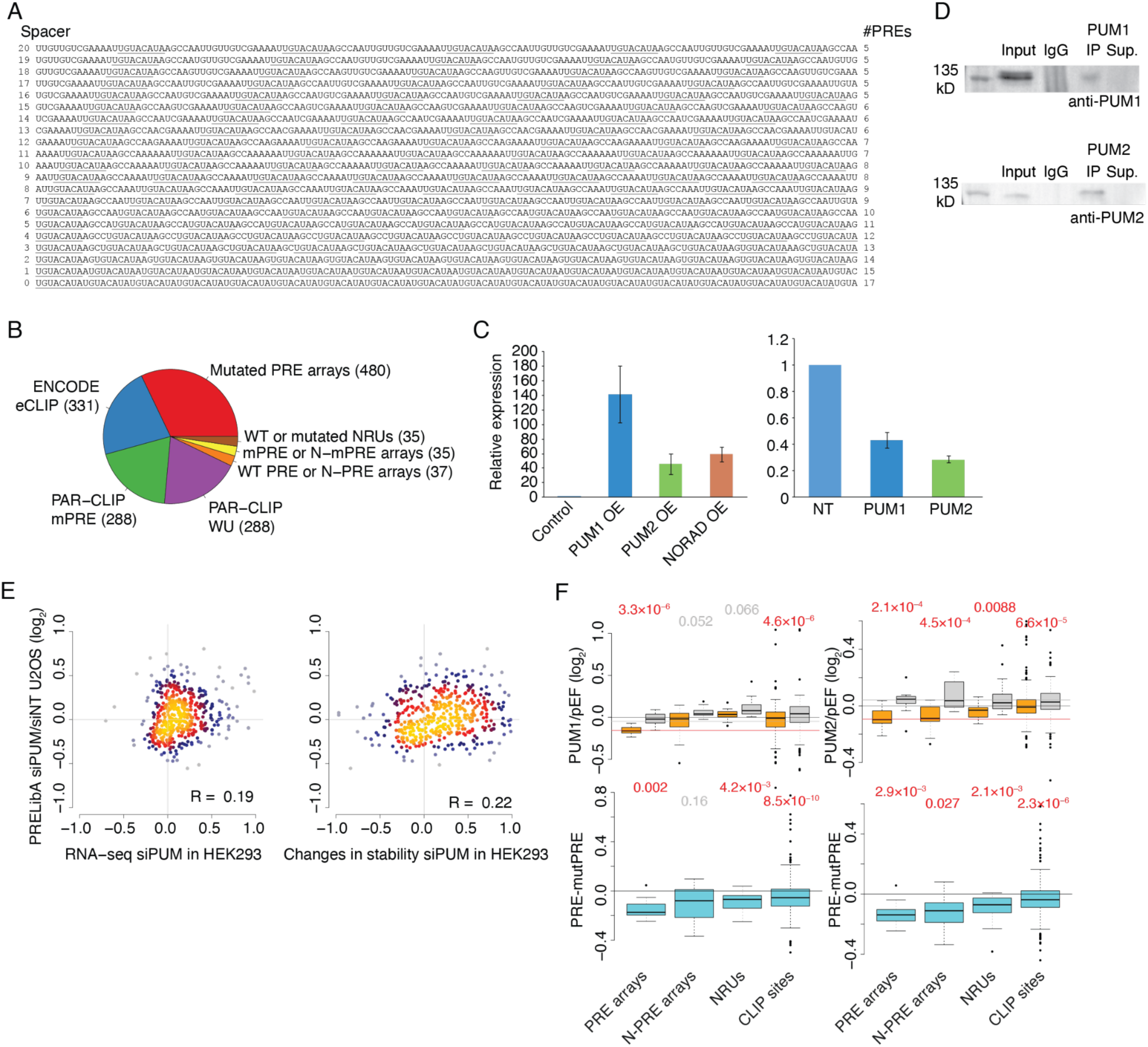
**(A)** Sequences of PRELibA sequences where the PREs (underlined) are separated by spacers with the indicated lengths. The eventual number of PREs in the sequence is shown on the right. **(B)** Composition of PRELibB, the numbers indicate the number of sequences in each category. **(C)** Left: qRT-PCR estimation of the changes in the expression of PUM1, PUM2, and NORAD in cells transfected with the indicated plasmid, all normalized to cells transfected with their corresponding empty plasmid (pcDNA3.1 for NORAD, pEF-BOS for PUM1/PUM2). Right: Changes in expression of PUM1 and PUM2 in cells transfected with siPUM (siRNAs for both PUM1 and PUM2), relative to an siNT transfection. **(D)** Western blot showing IP of PUM1 (top) and PUM2 (bottom) blotted with the corresponding antibody. **(E)** Correspondence between the changes in expression (left) and stability (right) of the endogenous genes in HEK293 cells following siPUM transfection (from (8, 15)) and the changes in expression of the fragments derived from these genes in our screen. **(F)** As in Fig. 2A for OE of PUM1 (left) and PUM2 (right), normalized to the empty vector pEF-BOS.

**Figure S2.**
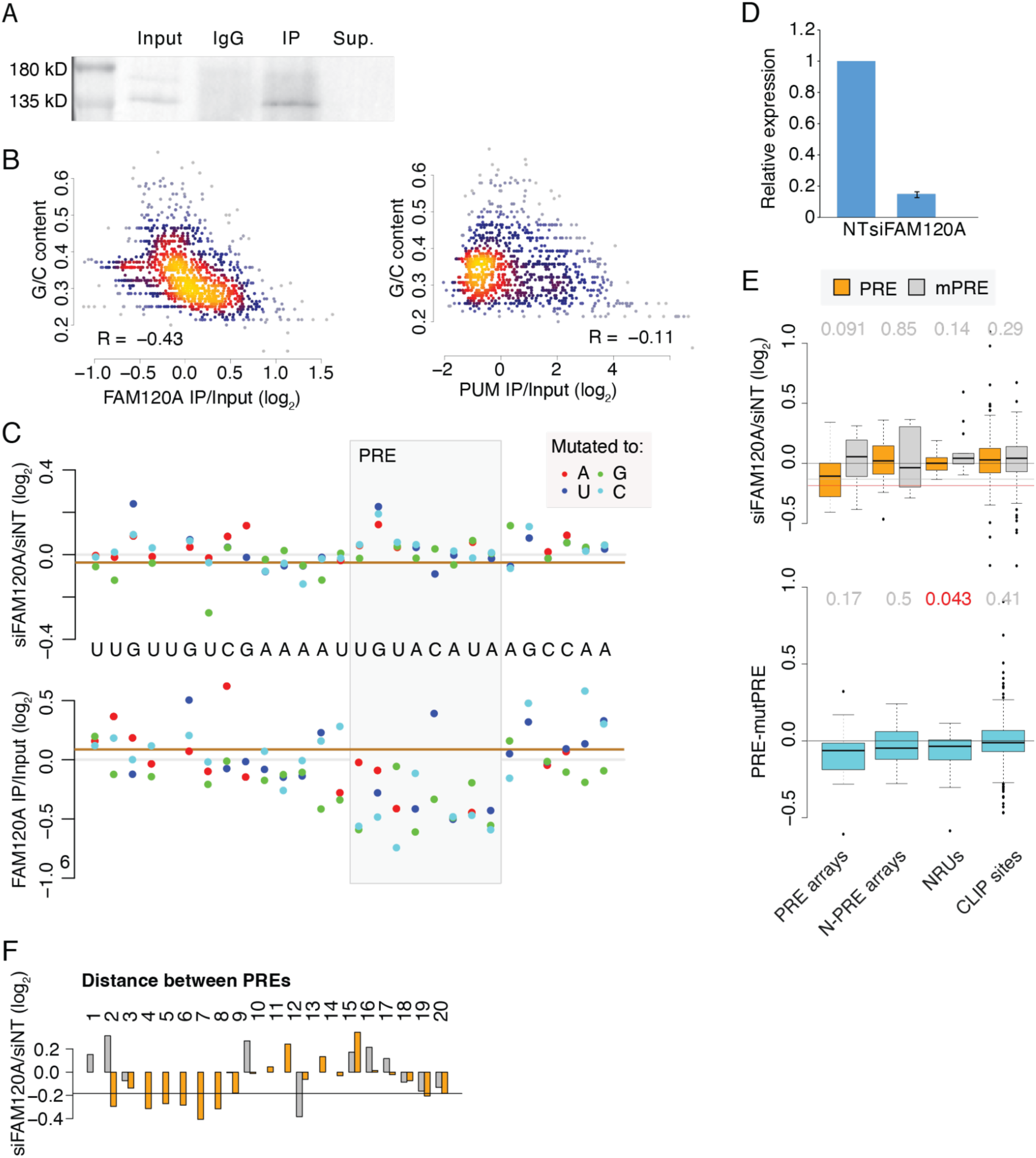
MPRNA and MPRNA-RIP for FAM120A. **(A)** Western blot for FAM120A for the indicated fractions. **(B)** Correspondence between binding of FAM120A (left) and PUM (right) and G/C content of the sequence. **(C)** As in Fig. 3D, showing the effects of mutations in the PRE arrays in response to siFAM120A and on FAM120A binding. **(D)** Changes in expression of the FAM120A mRNA following transfection of a nontargeting (NT) siRNA or the siFAM120A siRNA pool. **(E)** As in Fig. 6C, comparing the response to siFAM120A between subsets of PRELibA sequences. **(F)** As in Fig. 4A, comparing the response to siFAM120A between PRE- and mPRE-arrays with the indicated spacing between PREs.

**Figure S3.**
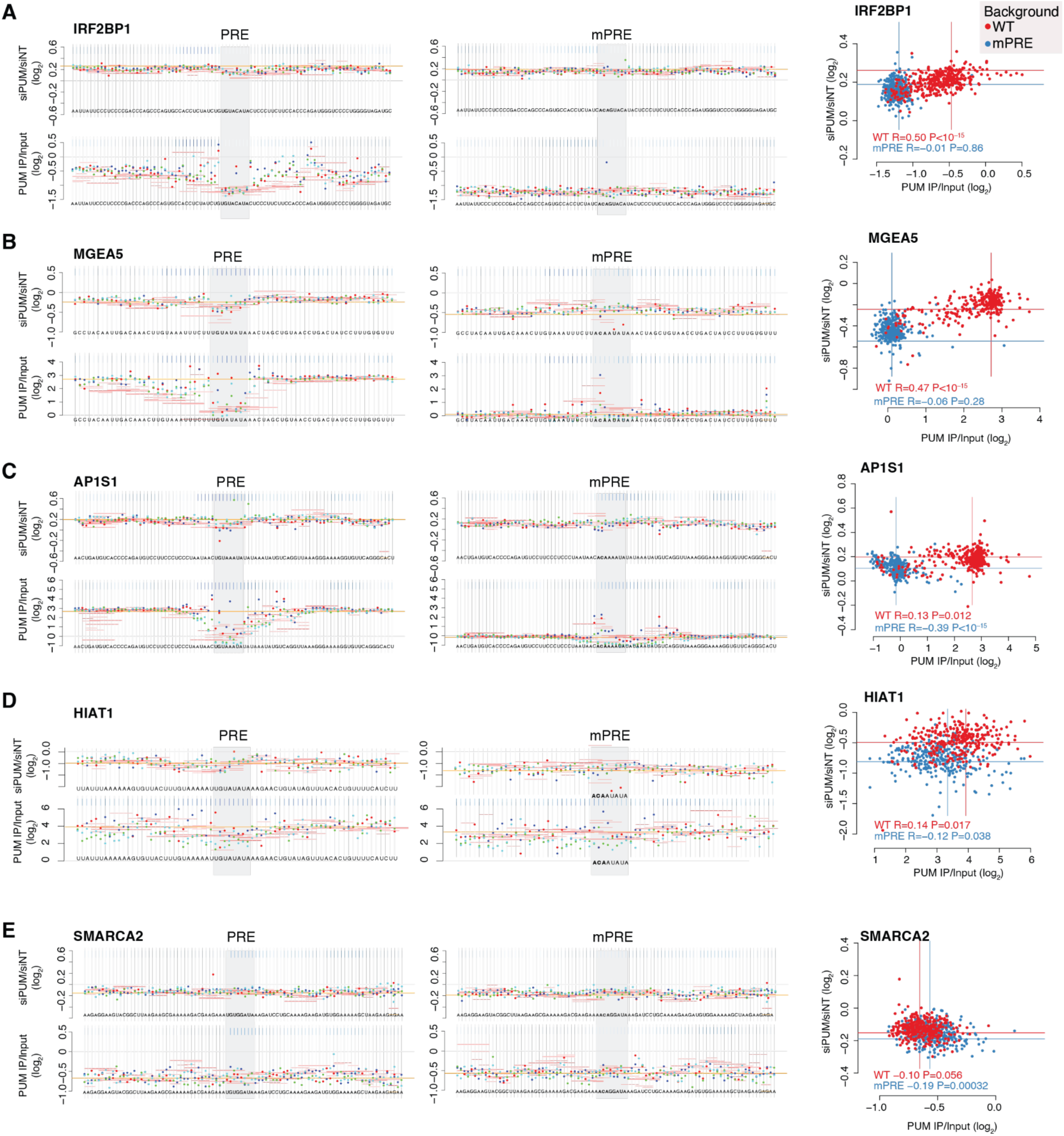
PRELibB sequences response to siPUM and binding to PUM. As in Fig. 7B, for the indicated sequences.

**Figure S4.**
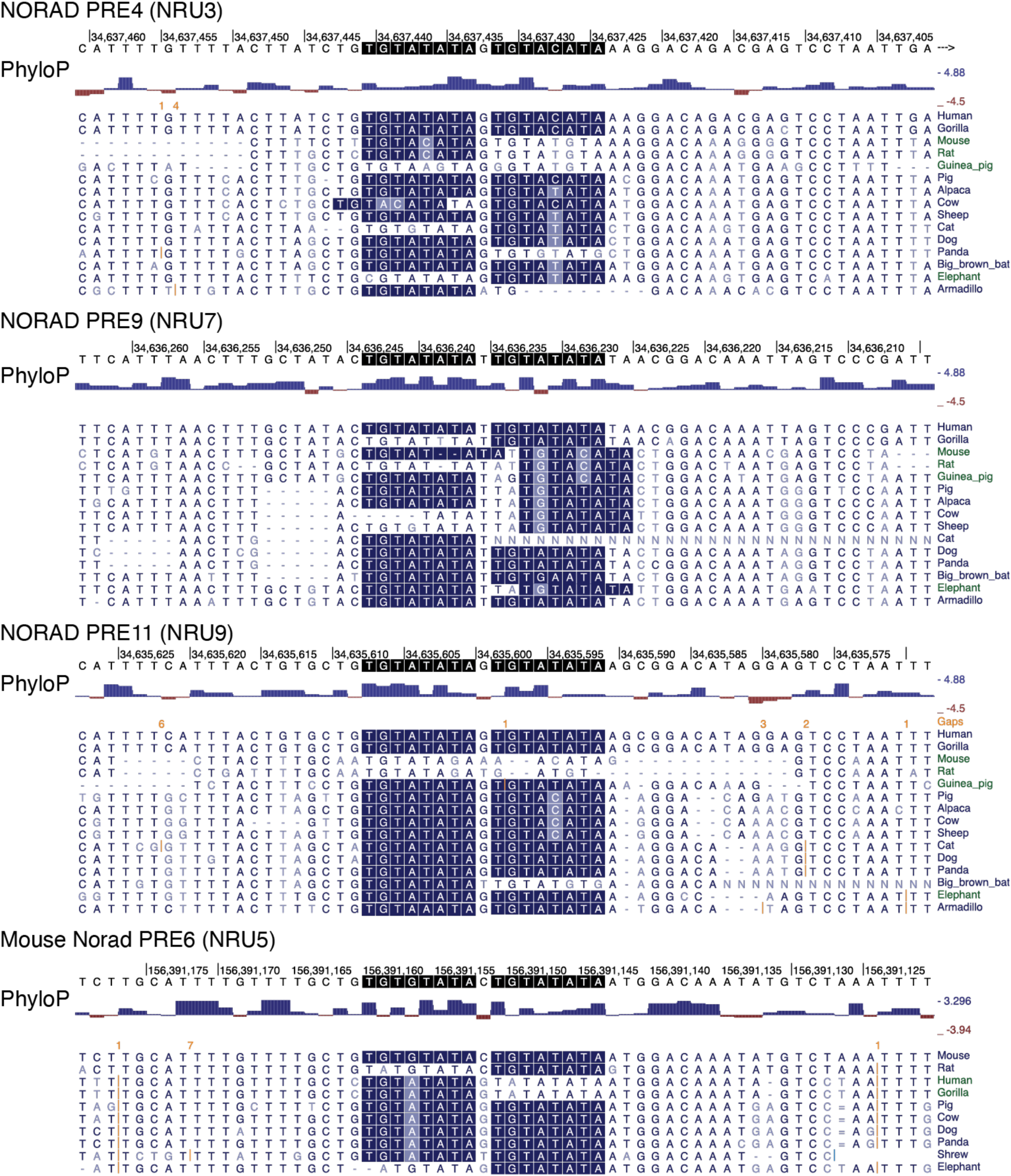
Conservation of tandem PREs in NRUs. Sequences and whole-genome alignments of regions in human *NORAD* (top three) and mouse *Norad* (bottom). PREs are highlighted.

## Supplementary Tables

**Table S1. Contents of PRELibA**

**Table S2. Expression levels, sequence features and DESeq2-computed ratios for PRELibA sequences.**

**Table S3. Contents of PRELibB**

**Table S4. Expression levels, sequence features and DESeq2-computed ratios for PRELibB sequences.**

**Table S5.**
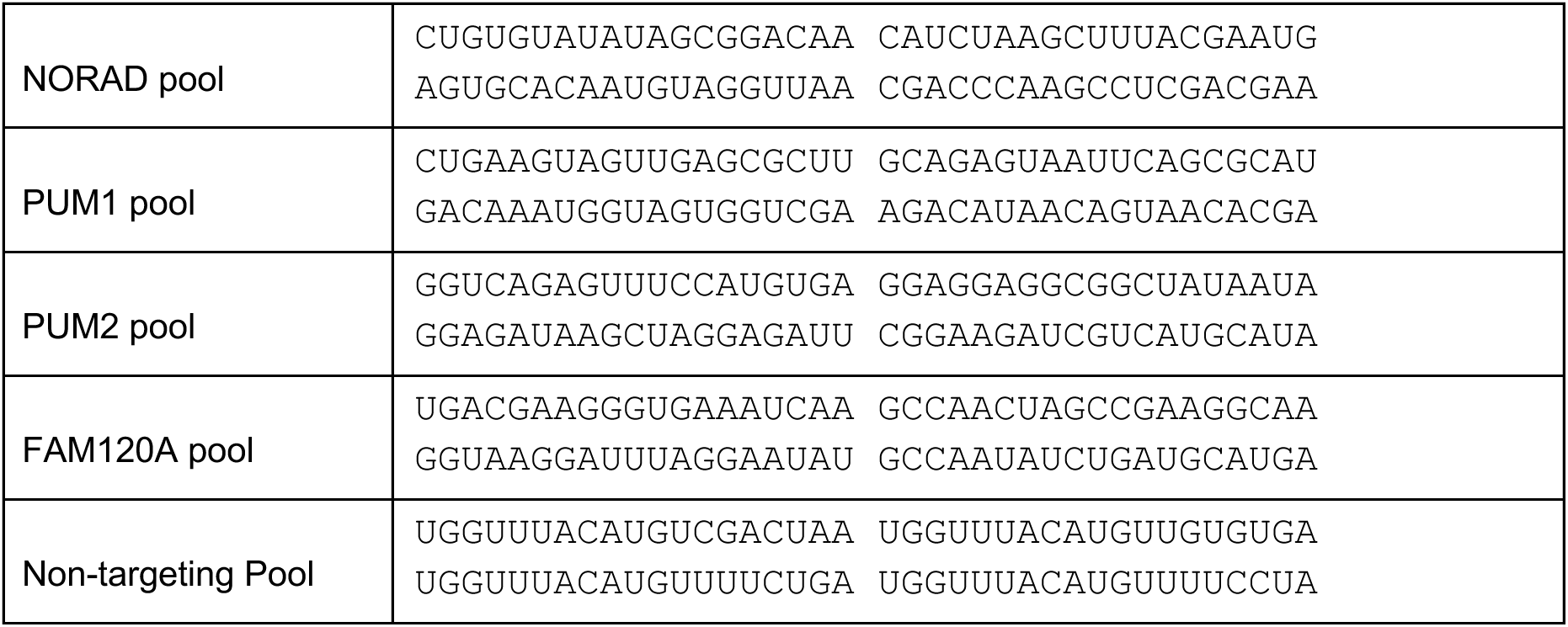
siRNA pool sequences.

**Table S6.**
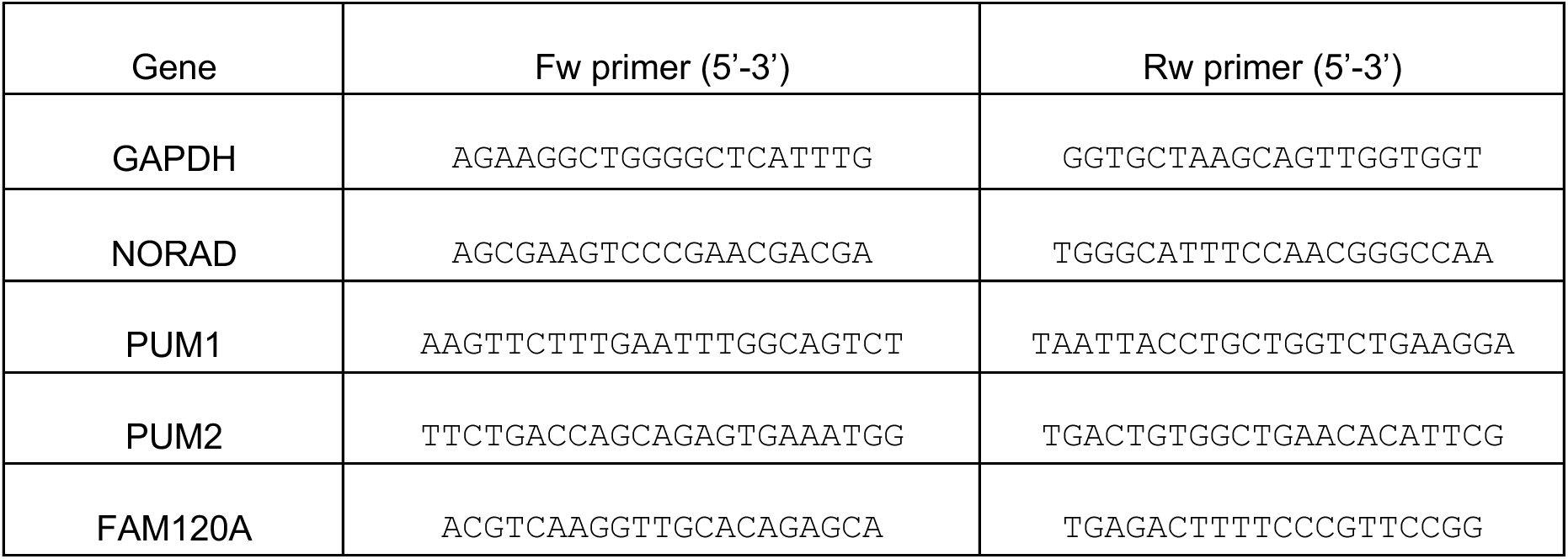
Primer sequences for Real-time PCR.

**Table S7.**
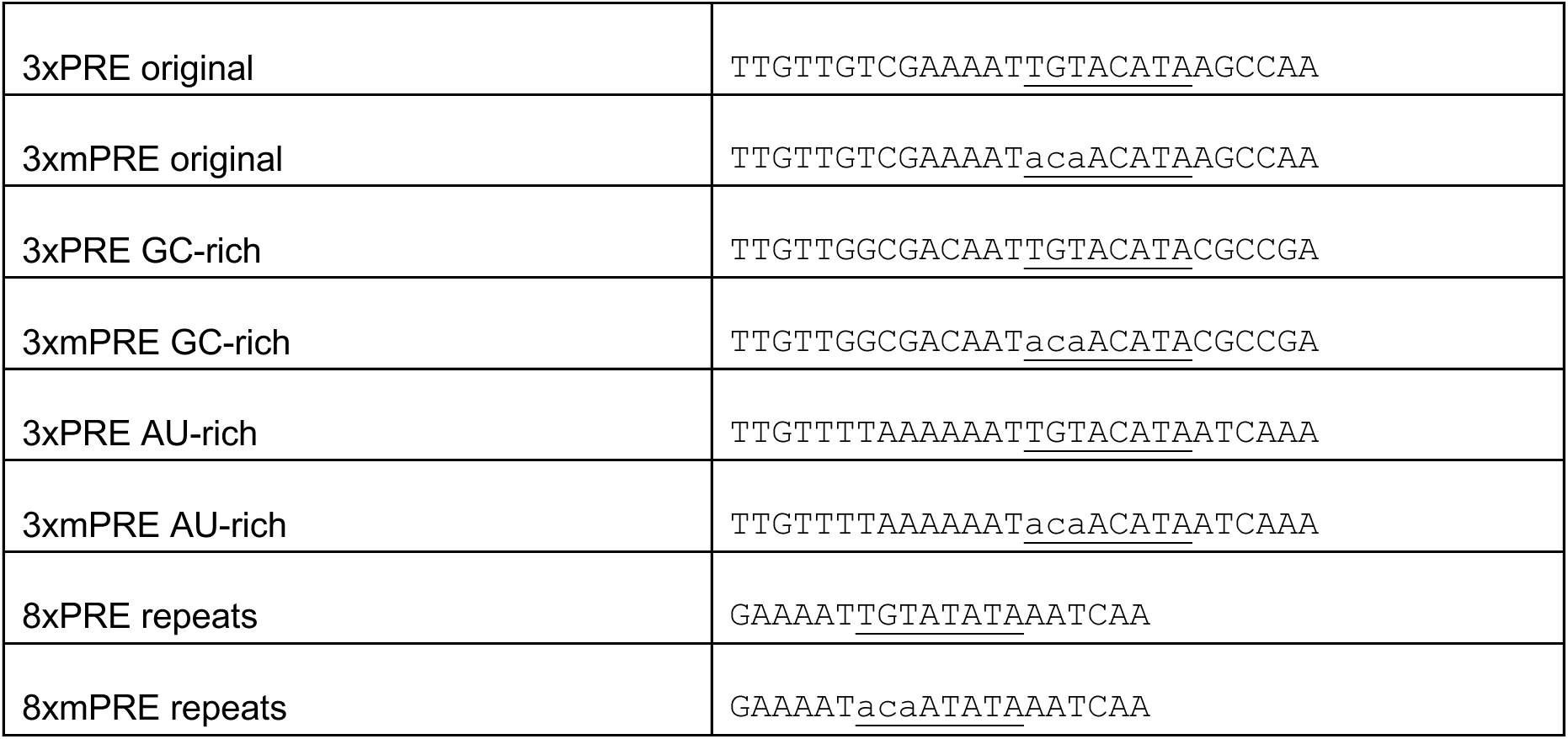
PRE and mPRE repeats used in plasmids.

